# Computer-guided enzyme engineering of PET hydrolase mutants towards improved PET affinity

**DOI:** 10.1101/2025.11.26.689686

**Authors:** Alexandra Balola, Sofia Ferreira, Caio Silva Souza, Diana Lousa, João Correia, Cláudio M. Soares, Isabel Rocha

## Abstract

Polyethylene Terephthalate (PET) is an extensively used plastic whose durability and resistance to degradation contribute to growing environmental pollution and concerns. Enzymatic PET degradation, particularly via PETase from *Ideonella sakaiensis*, has emerged as a sustainable approach due to its ability to depolymerize PET under mild conditions. While research has largely focused on enhancing the enzyme’s thermal stability through distal mutations, less attention has been given to active-site engineering aimed at directly improving catalytic efficiency. Here, we used an automated in silico protein engineering platform called Gene Discovery and Enzyme Engineering (GDEE), designed to systematically explore mutations at the active site. By leveraging FastPETase (FP) as scaffold, we perform a high throughput generation of thousands of variants, evaluated them via docking studies with a PET substrate analogue, and ranked candidates based on binding affinity and catalytic geometry. We identified S238Y as a key mutation that enhanced PET film degrading performance at 40 °C when inserted in two of the most active PETase variants reported to date: 2.2-fold increase in the FP scaffold and 3.4-increase in the ThermoStable-PETase (TSP) background. Compared to wild type PETase, FP S238Y showed a 14.8-fold increase in bulk activity, translating into 9.4-fold more TPA and 20-fold more MHET by UPLC, while TSP S238Y reached a 25.8-fold increase (14.4-fold more TPA and 42.6-fold more MHET). This mutation also enhanced catalytic efficiency and resistance to enzyme concentration inhibition, especially in the TSP scaffold. Molecular dynamics confirm position 238 as a relevant modulator of ligand stabilisation. These findings underscore the potential of targeted active-site engineering, combined with structure-guided prediction, to accelerate the development of efficient mesophilic biocatalysts for plastic waste remediation.

## INTRODUCTION

Polyethylene Terephthalate (PET) is a highly durable petroleum-derived thermoplastic, widely used in single-use packaging (e.g., bottles, containers, plastic bags), making it especially prone to mismanagement and a large contributor to pollution and ecosystem contamination ^1–4^. In 2023, 25.66 Mt of virgin PET were produced, yet conventional recycling remains insufficient: mechanical methods degrade polymer quality with each cycle, limiting reuse, while chemical depolymerization is energy-intensive, catalyst-dependent, and requires complex purification steps ^5–7^. Furthermore, despite advances in collection, sorting and recycling, most PET continues to be incinerated for energy recovery, landfilled or lost into the environment ^8,9^. In addition, when it reaches the environment, PET weathering generates microplastics—particles whose ecological and health impacts are only beginning to be understood—further underscoring the urgent need for innovative, circular-economy solutions ^1,10^.

PET is composed of two ester linked monomers: ethylene glycol (EG) and terephthalic acid (TPA) ^6^. Enzymatic PET depolymerization, where enzymes cleave this ester bond, currently stands out as a promising sustainable alternative to conventional recycling, with process-level analyses projecting significant energy savings, reduced greenhouse-gas emissions compared to fossil-derived PET production, and decreased toxicity issues ^11^. Since the first report of a PET-hydrolysing enzyme (PHE) two decades ago ^12^, numerous others have been identified and optimized ^13,14^. Among them, PETase stands out as the first, and so far, only enzyme known to have evolved specifically to hydrolyse PET as its primary and natural substrate. Since its discovery in 2016 from the bacterium *Ideonella sakaiensis,* PETase has become one of the most studied PET hydrolases ^15^. *I. sakaiensis* is able to use PET as energy and carbon source by secreting both PETase, which cleaves PET into bis(2-hydroxyethyl) terephthalate (BHET), mono(2-hydroxyethyl) terephthalate (MHET), and TPA, as well as MHETase that further hydrolyses MHET into TPA and EG. Unlike thermophilic PHEs which require elevated temperatures near PET’s glass transition temperature (Tg, 60–80 °C ^16^), PETase efficiently degrades PET at mesophilic temperatures ^15^, thereby avoiding the PET thermal aging and crystallization issues that hinder thermophilic hydrolases ^15,17,18^ PETase is a cutinase-like monomer with a canonical ɑ/β-hydrolase fold and a conserved Ser–His–Asp catalytic triad (S160–H237–D206) located on a shallow, hydrophobic cleft, considered central for binding of the large hydrophobic PET ^19,20^. Distinguishing it from most homologous enzymes, PETase has an extra disulfide bridge (C174–C210) that links and stabilizes loops containing catalytic residues H208 and D177 ^19,21^, and a flexible W185 residue near the catalytic site that may facilitate substrate positioning ^19,22^. Despite its potential, native PETase is limited by modest thermostability and catalytic efficiency; consequently, intensive improvement efforts through rational design ^23,24^, machine learning guided mutagenesis ^25,26^ and adaptive laboratory evolution ^27,28^ have yielded variants with markedly improved thermostability and activity. An early example is ThermoPETase (TP; S121E/D186H/R280A), rationally designed with observed increase the Tm from 48.81 °C in the wild type (WT) to 57.62 °C, and a boost of PET film degradation up to 14-fold after 72 h at 40 °C ^23^. ThermoStable-PETase (TSP) is derived by the incorporation of a disulfide bond N233C/S282C to TP, which considerably raised the Tₘ to 69.4 °C, boosted initial degradation rates by 20 % and increased product accumulation from PET films at 30°C ^24^. FastPETase (FP), also derived from TP through structure-guided machine learning, incorporates N233K and R224Q. Compared to TP, FP showed 2.4- and 38-fold higher activity at 40 °C and 50 °C, respectively, and outperformed other known hydrolases on untreated post-consumer PET at mild temperatures (30–50 °C), including DuraPETase ^25^ and the highly active thermophilic _LCC_ICCM 26.

So far, most PETase engineering efforts have come from amino-acid changes outside the catalytic cleft, such as disulfide bridges, salt bridges, and loop-stabilizing mutations that raise thermal stability and robustness and (consequently) boost activity, advancing their potential for industrial-scale applications. However, much less work has targeted residue modifications within the active site itself, even though tuned catalytic contacts could push PET breakdown even further. To explore that untapped space, we apply our in-house Gene Discovery and Enzyme Engineering (GDEE) pipeline ^29^ to redesign the catalytic site for higher efficiency. Our workflow (**Figure 1**) systematically introduces mutations at strategically selected active-site positions, generating thousands of enzyme variants. FP was chosen as the structural scaffold for its distal mutations that enhance thermostability without altering the catalytic site ^26^. By building on such stabilized variants, GDEE facilitates focused exploration of beneficial substitutions at the active site to increase activity further. Using this approach, we identified a key mutation S238Y that significantly improves PET film degradation in vitro across two different PETase variant scaffolds (both FP and TSP), improves initial degradation rates and decreases substrate-inhibition at high enzyme concentrations. This work highlights the potential of structure guided computational protein design to accelerate enzyme optimization.

**Figure 1.**
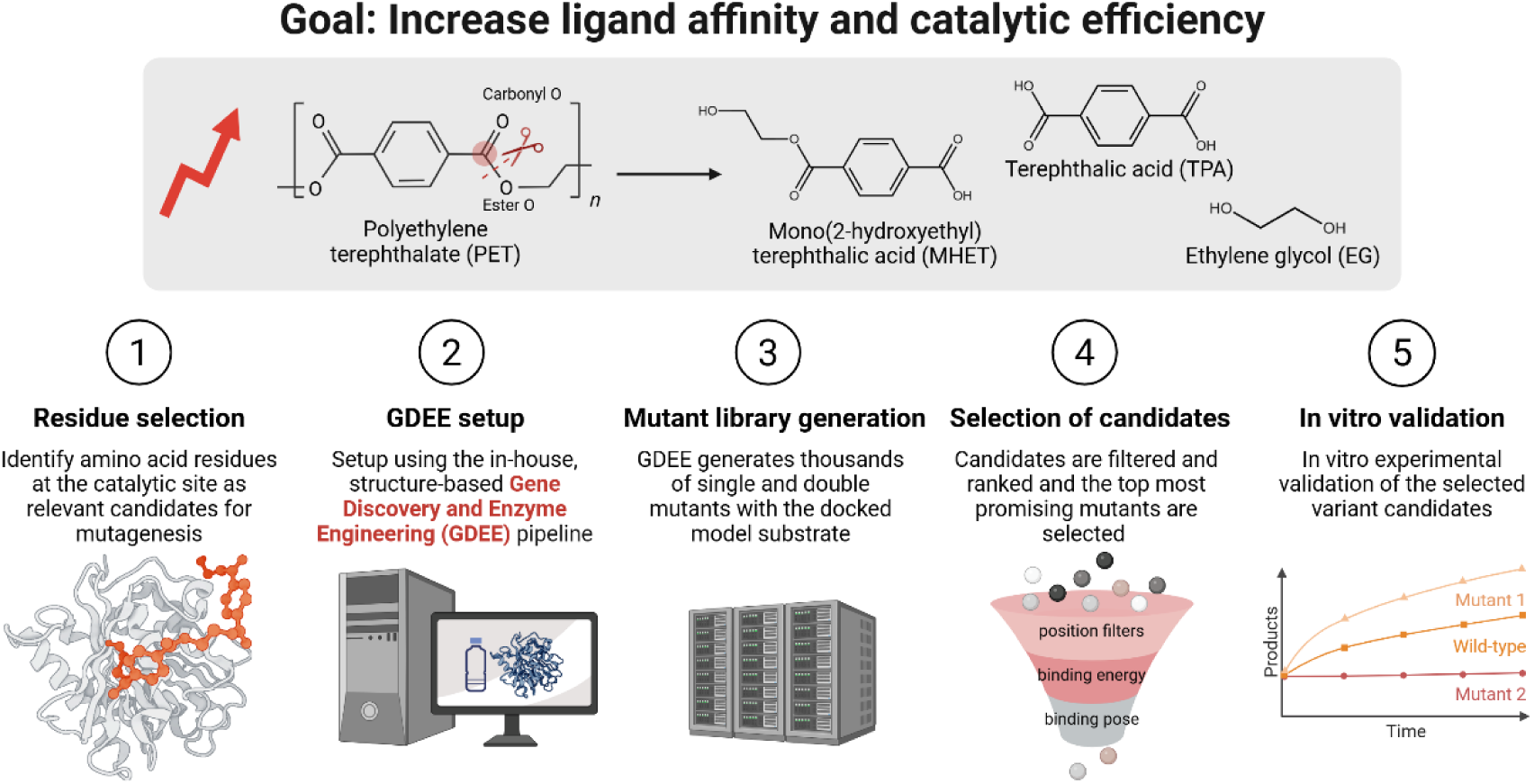
Scheme of the workflow used in the present study. A representation of the chemical reaction of PET hydrolysis is shown on the top panel. On the bottom panel, the protein engineering workflow used in this study is shown. Scheme created with BioRender.com.

## METHODS

### Section 1: In silico generation of candidate PETase variants

**Molecular Docking**: Molecular docking was performed using AutoDockTools (ADT) 1.5.6 ^30^ for input files preparation and AutoDock Vina 1.1.2 ^31^ for docking simulations. OpenBabel ^32^ was used for file format conversion, and PyMOL (The PyMOL Molecular Graphics System, Version 3.1.4.1 Schrödinger, LLC) was used for molecular visualization. The receptor structure FastPETase (PDB: 7SH6) was pre-processed by removing water, ions, and hydrogens. Missing atoms on residue T29 were added with PyMOL’s mutagenesis tool. Amino acid residues with alternative conformations were evaluated based on occupancy, retaining only the most occupied conformations. For position W185, alternative conformation B was selected. Polar hydrogens, Gasteiger charges and histidine protonation states (H104 and H186 as HE2; H237 as HD1) were added in ADT. The substrate 2-hydroxyethyl-(monohydroxyethyl terephthalate)2 (2HE-MHET2) was constructed in PyMOL and geometry was optimized in Avogadro2 (UFF, steepest descent, 2000 steps). The p*K*a of the hydroxyl sides were predicted as 12.8 with MolGpka ^33^, which were therefore considered protonated. Ligand torsions and charges were assigned with ADT and bonds were left rotable. Docking was targeted to the catalytic cavity using a 30 × 33 × 33 Å box, which ensured full coverage of the binding region and all possible ligand conformations. Docking was made with an exhaustiveness of 200 and generation of 50 docking poses for each ligand.

### Gene Discovery and Enzyme Engineering (GDEE) pipeline

Enzyme engineering of PETase was performed using the in-house, structure-based Gene Discovery and Enzyme Engineering (GDEE) automated pipeline ^29^, implemented as a Python package. Although this platform supports two main applications: gene discovery (where enzymes from online databases are screened to identify candidates structurally similar to a query protein for a given reaction) and enzyme engineering/optimization, this work will focus solely on the latter. In this mode, the GDEE pipeline leverages either an experimental or predicted enzyme structure and introduces, in a high-throughput manner, mutations at selected active-site residues to generate large variant libraries, which are then docked with the target ligand. Ligand binding affinities are used here as a proxy indicator for catalytic efficiency. The entire workflow is automated through modular Python scripts, which integrate required packages and software, and define parameters for each engineering round, including residue selection, mutation types (e.g., single and/or double mutants), distance metrics, and result filtering and ranking. All computations were run on an in-house high-performance cluster, using 20 nodes, each with 64 cores and equipped with four AMD Opteron^TM^ 6274 processors, enabling large-scale parallelization.

An automated engineering round by GDEE consists of: (1) mutagenesis, (2) modelling and quality assessment, (3) docking, and (4) ranking and filtering of the library poses. All structural parameters, docking parameters and specifications are configured prior to the start of the engineering round. Engineering was performed using the processed FP structure as described above.

Residues selected for mutagenesis were visually analysed for pocket geometry, proximity to catalytic residues, and charge to define permissible substitutions, excluding disruptive ones (e.g., glycine, proline, or bulky residues in constrained regions; **Table S1**). This rule set yielded a combinatorial space of 5,910 single and double mutants. For each mutant, local refinement within an 8 Å radius was performed using Modeller ^34^, generating 15 optimized models—88,650 in total. During structural optimization, the catalytic residues S237, H206, and D160 were kept immobile to preserve the active site. These models were ranked using two quality metrics (Modeller’s normalized DOPE [Discrete Optimized Protein Energy] score and the VoroMQA score ^35^) and the top five structures per mutant advanced to docking, reducing the library to 29,550 mutant structure models. Docking of each model was performed with Autodock Vina using the parametrized 2HE-MHET2 (box size: 30 × 33 × 33 Å; exhaustiveness: 200). Key interatomic distances defining catalytically productive binding poses and ligand orientations were pre-configured and automatically measured in every docking pose. Each docking simulation produced 500 binding poses per model, resulting in 14,775,000 docking poses, which were filtered using the distance metrics (**Figure S1**). Remaining variants were ranked by predicted binding free energy. Top candidates were visually inspected, and promising mutants were selected for in vitro validation, excluding those previously reported in the literature as unproductive.

### Multiple Sequence Alignment

Putative PET hydrolase sequences were retrieved from the PET-PAZy (plastic-active enzymes) database (accessed on 09/05/2025), when available ^13^. To ensure biological relevance, sequences showing less than 45% identity to PETase were excluded based on pairwise identity calculated using the online BlastP tool (**Table S2**). All selected sequences were aligned using T-Coffee Alignment Server (version 11.00) with default parameters, and the amino acid frequencies in selected positions were extracted.

### Section 2: Molecular biology and protein production

#### DNA sequences and cloning

The PETase sequence from *I. sakaiensis* (WT; GenBank: GAP38373.1) was codon-optimized for expression in *Escherichia coli* K12 using the IDT Codon Optimization Tool, and synthetized by IDT (Iowa, USA). Mutations S121E/D186H/R280A (TP), S121E/D186H/R280A/R224Q/N233K (FP) and S121E/D186H/R280A/N233C/S282C (TSP) were introduced into the codon-optimized WT sequence of PETase, and the resulting sequences were synthetized by GeneCust (Boynes, France). The native signal peptide was predicted using the SignalP 5.0 server ^36^, identifying residues 1–27 as the PETase signal peptide, which was removed during cloning experiments. All PETase sequences were individually cloned into the pETDuet-1 plasmid from the Novagen pET System (Merck Millipore, Massachusetts, United States) with an N-terminal His-tag for purification. Primers were designed using the NEBuilder® v2.7.1 (New England Biolabs) tool (**Table S3**) and synthesized by IDT (Iowa, USA). Cloning was carried out using the FastCloning protocol ^37^, with polymerase chain reaction (PCR) amplification performed using Phusion High-Fidelity DNA Polymerase (ThermoScientific, Waltham, United States) according to the manufacturer’s instructions. Constructs were transformed by heat-shock into chemically competent *E. coli* DH5α cells (New England Biolabs) and colonies were picked for colony PCR. Briefly, each colony was resuspended in 20 µL Milli-Q H2O, and 2 µL of this suspension was used as the PCR template. Amplification was performed with DreamTaq DNA Polymerase (ThermoScientific, Waltham, USA) using primers DuetUp1 ad and DuetDown1 (**Table S2**), following the manufacturer’s instructions. Plasmids were extracted using the ZRPlasmid Miniprep-Classic Kit (Zymo Research) and sequence-verified by Sanger sequencing (StabVida, Lisbon, Portugal).

#### Site-directed mutagenesis

Mutant gene sequences were generated by site-directed mutagenesis using a two-step PCR protocol with Phusion High-Fidelity DNA Polymerase (ThermoScientific) and mutation-specific primers designed with the QuickChange® Primer Design tool ^38^ (Agilent), synthesized by IDT (Iowa, USA) (**Table S4**). The protocol, adapted from Reikofski and Tao (1992) ^39^, involved two steps: first, two separate single-primer PCR reactions were carried out for 5 cycles. In the second step, 25 μL of each reaction was combined, mixed with 1 U of Phusion polymerase, and further amplified for 30 cycles. PCR products were digested with 1 μL *DpnI* (ThermoScientific) for 1 h at 37 °C, and 6–8 μL of the digest were transformed into *E. coli* DH5α (New England Biolabs). Constructs carrying the desired mutations were confirmed by Sanger sequencing (StabVida) and used as templates for further mutagenesis when needed.

#### Bacterial strains

For enzyme expression, correctly assembled plasmids were transformed into the protein production strain *E. coli* BL21 (DE3) (NZYtech, Portugal) using heat-shock transformation into chemically competent cells. All strains used in this study are summarized in **Table S5**.

#### Protein purification

Production strains from **Table S5** were cultured in 1 L Luria-Bertani (LB) medium with 50 µg mL^−1^ ampicillin, at 37 °C and 150 rpm. At OD600 of 0.4–0.5, protein expression was induced with 0.1 mM isopropyl β-D-1-thiogalactopyranoside (IPTG), and cultures were incubated overnight at 16 °C, 150 rpm. Cells were harvested by centrifugation at 8,600 x g (Beckman JA-10 rotor) for 10 min, resuspended in 15 mL of Buffer A (20 mM potassium phosphate pH 7.5, 500 mM NaCl, 40 mM imidazole) containing DNase I and EDTA-free protease inhibitor, and lysed with the French Press. Lysates were clarified by centrifugation at 48,200 x g (Beckman JA-25.50 rotor) for 1 hour at 4 °C, filtered through a 0.2 µm membrane, and purified on a 5 mL HisTrap HF Ni-NTA column (GE Healthcare) using an ÄKTA Purifier 10 chromatography system (GE Healthcare). Elution was performed using Buffer B (20 mM potassium phosphate pH 7.5, 500 mM NaCl, 500 mM imidazole) over 20 column volumes at 5 mL min^−1^, and 5 mL fractions were analysed by SDS-PAGE. Protein-containing fractions were pooled, concentrated to 10–15 mL, and buffer-exchanged into 20 mM potassium phosphate, 500 mM NaCl using a 10 kDa Molecular weight cutoff (MWCO) Vivaspin concentrator. Protein concentration was determined via absorbance at 280 nm using Nanodrop OneC (ThermoFisher), and samples were aliquoted and stored at −80 °C.

### Section 3: In vitro assays

#### PET films

Amorphous PET films (ES301445; 250 µm thickness) were purchased from Goodfellow Ltd. (Germany). This PET film has a crystallinity of 2–2.3 % ^17,40^. BHET (B3429) and TPA (T0166) were purchased from TCI Europe N.V., while MHET (BD00777875) was purchased from BLD Pharmatech Ltd.

#### In vitro enzymatic assays

Enzymatic assays were adapted from Zhong-Johnson et al. ^24^ with minor modifications. Reactions (1 mL) were performed in 16 mm borosilicate tubes (DWK Life Sciences), containing 980–960 µL reaction buffer (50 mM glycine-NaOH pH 9.0, 50 mM NaCl, 10 % DMSO [v/v]) and 20 or 40 µL of PETase stock (7.5 µM), to achieve a final concentration of 150 nM or 300 nM. A 6 mm amorphous PET disk was submerged in the reaction medium and tubes were sealed with parafilm. Incubation was carried out at 40 °C for 72 h at 200 rpm. Absorbance at 260 nm was measured at 24, 48, and 72 h using 4 µL aliquots on a NanoDrop OneC. A no-enzyme control was included in all assays. After 72 h, the PET disks were removed, reactions were heat-inactivated (85 °C, 20 min), centrifuged (18,400 x g on an Eppendorf FA-24x2 rotor, 10 min), and supernatants were stored at 4 °C for analysis. Calibration curves were obtained by measuring TPA and MHET standards (5–200 mg L^−1^) at 260 nm using a NanoDrop OneC. Linear regressions gave extinction coefficients of 5464 M^−1^ cm^−1^ for MHET and 4233 M^−1^ cm^−1^ for TPA (correlation coefficient [R^2^] = 0.99 for both), in agreement with literature values ^24,41^. The MHET coefficient was used to estimate bulk product concentrations as MHET equivalents (MHETEq) following the Beer–Lambert law. Each mutant was tested in 3–5 independent biological replicates with two technical replicates. Outlier analysis (IQR method and Grubbs’ test) was performed at both the technical and biological levels. For analysis, technical replicates were averaged to represent each biological replicate.

#### Enzymatic evaluation at different pH

To test activity under different pH conditions, additional buffers (50 mM Tris-HCl, 50 mM NaCl, 10% DMSO [v/v]) were prepared at pH 8.0 and pH 7.5 and used in an otherwise equal in vitro enzymatic assay setup at 40 °C.

#### Activity at different enzyme concentrations

Stock solutions of each PETase variant (2.5, 7.5, and 70 µM) were prepared to achieve final concentrations of 25–400 nM by adding 20 or 40 µL to 1 mL reactions. Absorbance at 260 nm was measured at 1, 2, 3, and 4 h, using 1.5 µL aliquots on a NanoDrop OneC, to calculate initial reaction rates, and at 24, 48, and 72 h to assess long-term product accumulation. Absorbance values were plotted over time for each enzyme concentration, and the slope of the early time points was used to calculate the reaction rate. Percentage inhibition was calculated as the ratio between the highest activity across all concentrations and the activity at the highest concentration tested. Two biological replicates were performed per condition, following the same 40 °C assay protocol described above. Kinetic parameters were derived by nonlinear regression of the initial reaction velocities fitted to the inverse Michaelis-Menten model equation (**Equation 1**).

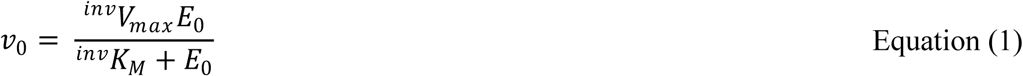

#### Analytical analysis

Quantitative analysis was performed using an ACQUITY Ultra Performance Liquid Chromatograph (UPLC) system (Waters, Massachusetts, USA) equipped with a photodiode array (PDA) detector and a Symmetry C18 reverse-phase column (5 μm, 4.6 mm × 250 mm, 100 Å pore size). Separation was carried out using a binary gradient of Buffer A (water with 0.1 % formic acid) and Buffer B (100 % acetonitrile), with the following gradient profile: 85:15 (A:B) at 0 min, 50:50 at 20 min, returning to 85:15 at 22 min and held until 30 min. The flow rate was maintained at 0.8 mL min^−1^, with the column temperature set at 30 °C and sample temperature at 20 °C. Both samples and standards were injected with a volume of 7.5 µL, and analytes were detected at 260 nm. Calibration curves for TPA, MHET, and BHET were prepared using standard solutions dissolved in 100 % DMSO across a concentration range of 1.0–200 mg L^−1^, using a minimum of eight calibration levels per analyte (R^2^ ≥ 0.99). BHET levels were not detected or were negligible (< 0.03 mM) in the reaction samples and were not included in the product sum.

### Section 4: Molecular Dynamics simulations

#### Ligand parametrization

Antechamber ^42^ was used to create a previously gas-phase geometry optimised at the HF/6-31G* level structure into a Gaussian 09 ^43^ input to generate electrostatic potentials in space to calculate partial atomic Electrostatic Potential (ESP) charges. ^43^ These potentials in space served as the target for a two-stage restrained ESP (RESP) fit ^44^. The RESP-charged MOL2 file with atom types taken from the General AMBER Force Field (GAFF2) ^45^, and parmchk2 ^42^ was used to identify and generate any missing force field parameters. AMBER’s tleap ^46^ combined the RESP-charged MOL2 and any supplementary parameter files from GAFF2 to produce an AMBER topology and coordinate files and a hydrogen-complete reference PDB structure, which were then converted to GROMACS format via ACPYPE ^47^. These were used to generate a ligand topology include file (.itp), for incorporation into subsequent molecular dynamics (MD) system assemblies.

#### Ligand-pose extraction and coordinate preparation

To transfer the docking-derived ligand pose into an MD-ready format, the GDEE-ranked poses were loaded in PyMOL, and the ligand for the selected mutant was exported as a PDB file. PyMOL was used to fix bond orders and add hydrogens to give an all-atom PDB. Python was used to map the coordinates from the docking pose onto the pre-parameterised AMBER ligand: atom names were matched one-to-one, coordinates overwritten, and a hybrid PDB retaining GAFF2 labels but adopting the docked geometry was produced. This file was converted to GROMACS format with editconf for MD simulations.

#### Protein preparation

Protein topology was generated from the engineered variant exported from the GDEE results with PyMOL (**Section 1**). Crystallographic waters from PDB 7SH6 were reintroduced and the missing native C-terminal S290 was added using PyMOL’s Builder; two waters molecules clashing with S290 were removed.

#### Protein-ligand system construction and minimization

All system preparation steps and MD simulations were performed with GROMACS 2021.7 ^48^ using the AMBER ff14SB force field ^49^ for the protein. Ligand poses were loaded into PyMOL along with the respective protein variant and water molecules within 3 Å of the ligand were removed. The resulting protein PDB was processed with GROMACS pdb2gmx to assign histidine protonation states (**Section 1**) and add terminal charges, yielding the initial GROMACS coordinate (.gro) and topology files. Ligand coordinates were merged into the protein coordinate file. The complex was placed in a dodecahedral periodic box with a 1.5 nm buffer, solvated, and neutralized with counterions to obtain a 0.1 M ionic strength. Energy minimization was performed with steepest descent for up to 50,000 steps, with a 10,000 kJ mol^−1^ nm^−2^ harmonic restraint on all heavy atoms. Electrostatics were treated with Particle-Mesh Ewald (PME, 0.8 nm real-space cutoff, Fourier grid spacing 0.12 nm), van der Waals interactions used a 0.8 nm cutoff with analytical long-range dispersion corrections, the Verlet neighbor-list scheme was employed and LINCS was used to constraint H-bond lengths. The time step used for the integration of equations of motion with the leap-frog integrator was 2 fs.

#### Initialisation and production

Initialisation comprised four consecutive 100 ps segments applying positional restraints to protein heavy atoms and ligand ring atoms. In the first, velocities were generated at 300 K and temperature was held at 300 K with a Berendsen thermostat (τ = 0.01 ps); in the second, the same thermostat was retained with τ increased to 0.1 ps. The third segment activated isotropic pressure coupling via a Berendsen barostat (τ = 1 ps, compressibility = 4.5×10^−5^ bar^−1^) at 1 bar. Finally, thermal coupling was switched to a V-rescale thermostat and pressure coupling to a Parrinello-Rahman barostat (τ = 2 ps, compressibility = 4.5×10^−5^ bar^−1^) at 1 bar. A unique random seed was generated at the start of each initialization to ensure independent, randomized initial velocity assignments. A 100 ns slow-release stage was then conducted in two consecutive 50 ns segments: the first with restraints on protein heavy atoms and ligand rings, and the second with restraints on Cα atoms and ligand rings. Unconstrained production runs of 200 ns were executed initiated from the last slow-release frame. Five independent replicate initialisation/production simulations were carried out for each mutant. All simulations were performed on a local workstation equipped with an AMD Ryzen 9 7950X 16-core processor and an NVIDIA GeForce RTX 4070 Ti GPU.

#### Trajectory analysis

MDAnalysis (version 2.9.0) ^50,51^ was used to manipulate, analyse and extract structural data from the MD data following standard analysis workflows. For each replicate simulation, the mean trajectory-derived structural metrics over the full production trajectory was calculated (n=5). These were used to estimate the uncertainty in the overall means for each variant using non-parametric bootstrapping with 1000 resampling iterations. The 95 % confidence intervals for the mean assuming normality, corresponding to ± 2 standard deviations, were used as error bars in the plots. The relationship between mean trajectory-derived structural metrics (dependent variable) and in vitro activity (independent variable) was assessed using simple linear regression, using SciPy’s linregress function ^52^, which returns the slope, intercept, Pearson correlation coefficient (*r*) and the *p*-value (*p*) for the correlation. Pearson *r* was calculated as the covariance of x and y divided by the product of their standard deviations. Extracted data was organized using pandas (version 2.2.3) ^53^ and plots were generated using Matplotlib (version 3.10.3) ^54^.

#### Root mean square deviation (RMSD)

RMSD was calculated for all Cα atoms with backbone alignment and for Cα atoms at the **active site cleft selection** (residues within ∼ 4 Å of the ligand poses: residues 86–93, 116–119, 159–161, 185–189, 204–214, 233–245, 280) with backbone alignment or active site cleft fitting to the first frame of each trajectory.

#### Ligand-binding cleft contacts

For each residue within the active site cleft selection, the percentage of simulation frames in which it was in contact with the ligand was calculated. A contact was defined when the distance of any pair of heavy atoms between the residue and the ligand is ≤ 4 Å, as determined using the distance_array function. To focus on catalytically relevant poses, only simulation frames in which the ligand was positioned within the catalytic groove were considered. This was defined as having at least one of the ligand’s catalytic carbon atoms ≤ 6 Å of the Oγ atom of residue S160.Residue-specific contact frequencies were used to write into the B-factor field of a representative MD simulation PDB file for visualization.

#### Hydrogen bond analysis

Protein-ligand hydrogen bonds were quantified for each run using MDAnalysis’ HydrogenBondAnalysis default geometric criteria (hydrogen-acceptor distance ≤ 3.0 Å and donor-hydrogen-acceptor angle ≥ 150°). Donor atoms were limited to residues within the active-site cleft, while acceptor atoms were restricted to the ligand. Both hydrogen and acceptor atoms were identified using MDAnalysis’ built-in guesser. Within each frame, multiple atom-level hydrogen bonds between the same donor residue and the ligand residue were collapsed to a single event to avoid double counting due to bifurcated interactions. Frames used for occupancy calculations were filtered by a proximity cutoff that defined the ligand as within the catalytic groove (≤ 6 Å between any of the ligand’s catalytic carbons and S160’s Oγ) and computed as the fraction of these hydrogen-bonded frames among the bound frames (≤ 6 Å set).

#### π stacking analysis

To analyse the aromatic stacking interactions between the ligands’ phenyl rings and the aromatic side chain of residues 185 and 238, the geometric centroids of each ring were computed for every simulation frame, and plane normal vectors were obtained from the cross product of two non-collinear bond vectors within each ring. The Euclidean centroid-centroid distance, normal-normal angle, and in-plane offset distance were calculated for each protein-ligand ring pair. Interactions were classified using literature-derived thresholds ^55–57^: centroid distance ≤ 5.0 Å for parallel interaction and ≤ 6.0 Å for T-shaped interactions, normal-normal angle for face-to-face interactions ≤ 30° and an in-plane offset ≤ 1.5 Å, while in-plane offset ≥ 1.5 Å was considered parallel-displaced; T-shaped interactions were defined by plane angles between 60° and 90°. For aromatic interaction total counts, only frames where the ligand was within the catalytic groove (≤ 6 Å between any of the ligand’s catalytic carbons) were considered.

## RESULTS

### Mutant selection through high-throughput generation and filtering of docking poses

To identify the most favourable binding conformations and select relevant residues at the binding cleft as candidates for mutation, molecular docking simulations were performed using FP (PDB: 7SH6) as the receptor and 2HE-MHET2 as a model PET substrate (**Figure 2A**). This ligand was chosen to avoid the incorrect binding of longer segments such as 2HE-MHET4 ^17,20^ or 2HE-MHET3, which in our tests folded back on themselves and yielded artificially high binding energies, not mimicking the polymer nature of the natural substrate. We allowed its bonds to remain rotatable during docking to better reflect the dynamic nature of PET chains: π-electron delocalisation and steric effects may favour a plantar TPA unit ^58^ but this tendency is not strong enough to justify fixing its aryl-alkyl ester bonds. Likewise, the EG dihedral naturally partitions between gauche (± 70–80°) and trans (180°) states, with amorphous PET favouring gauche (∼91:9 ratio) ^58,59^. Water-driven plasticization further lowers already lower surface-level Tg by ∼15 °C, increasing chain mobility at the liquid interface ^60–62^. Moreover, we retained the alternative conformation B for residue W185, which widens one end of the cleft and yielded more low-energy, productive poses. This rotamer, generally referred to as conformer “B”, reflects W185’s unique conformational mobility that broadens the cleft and better accommodates PET during binding ^19,22^.

**Figure 2.**
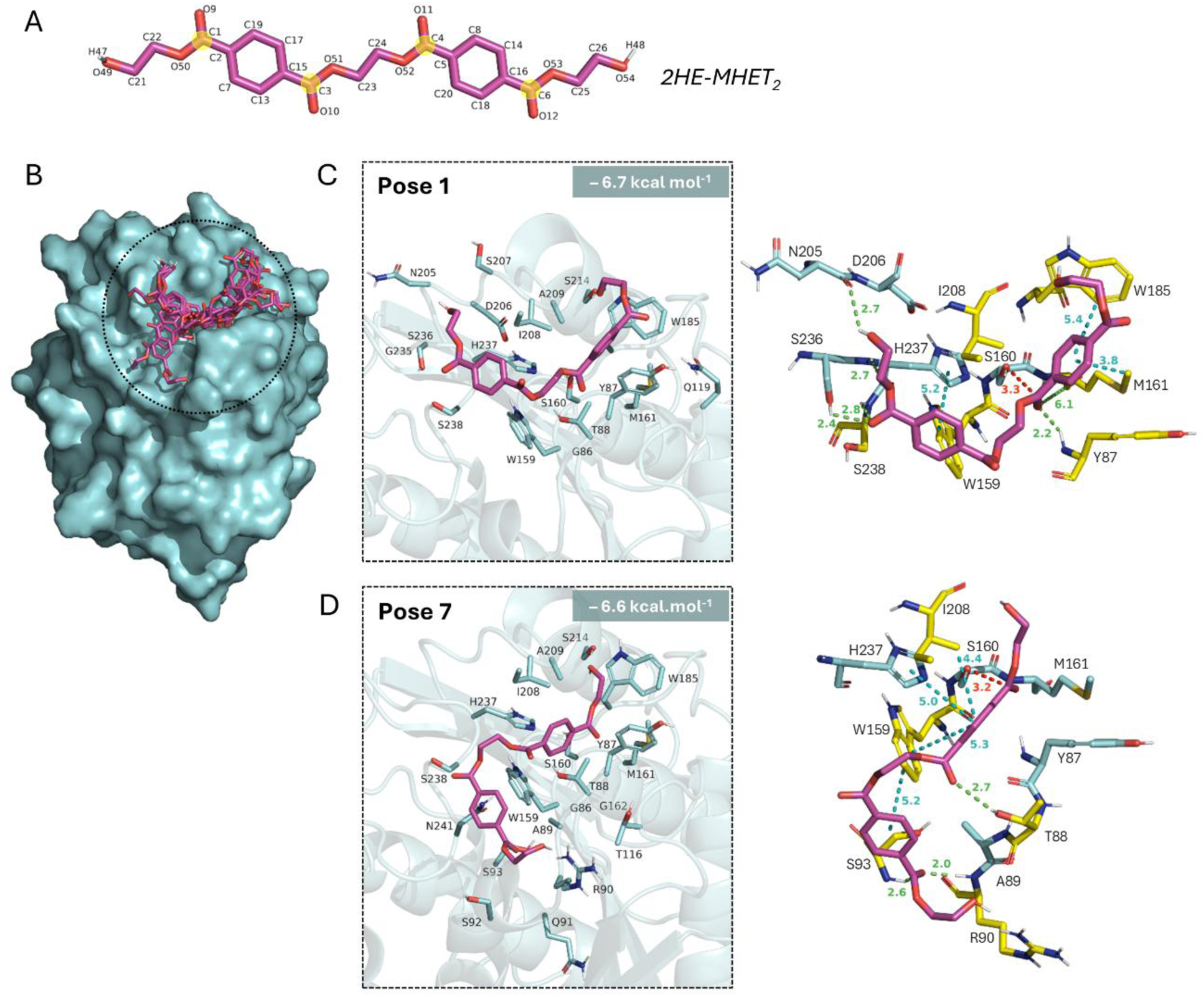
Docking poses and interaction networks of the ligand 2HE-MHET2 within the catalytic groove of FastPETase (FP; PDB: 7SH6). **(A)** Chemical structure of 2HE-MHET2, with the number of each atom represented and with the reactive carbonyl carbon atoms overlayed with a yellow circle. **(B)** Surface representation of the top 10 binding poses from docking calculations. **(C, D)** Cartoon and stick representations of the binding pocket showing the ligand (magenta) orientation for Pose 1 (C) and Pose 7 (D), with binding affinity energies of −6.7 and −6.6 kcal mol^−1^, respectively. On the left side, residues within 5 Å of the ligand are shown. On the right, close-up view of the molecular interactions stabilizing Pose 1 and Pose 7 are shown. Hydrogen bonds are shown as green dashed lines and aromatic interactions are shown in cyan. Distances are shown in angstroms (Å) and residues selected for protein engineering for each pose are shown as yellow sticks.

Docking results revealed a network of interactions stabilizing the ligand, which converged on the catalytic pocket (**Figure 2B**) and adopted two main favoured conformations, represented by Pose 1 (**Figure 2C**) and Pose 7 (**Figure 2D**), with binding affinities of −6.7 and −6.6 kcal mol^−1^, respectively. In Pose 1, a ligand’s reactive carbonyl sits 3.3 Å from S160, anchored for nucleophilic attack by a network of hydrogen bonds: backbone N-H groups of H237 and S238 interact with ester and carbonyl oxygens, respectively, while S236 adds a third interaction to the same oxygen contacted by S238. H237’s imidazole stacks π–π T-shaped with the first aromatic ring. Aromatic interactions tighten the complex: W185 and M161 engage the second ring through π–π T-stacking and π–σ interactions, and W159 provides hydrophobic stabilization at the cleft’s base. Pose 7 resembles Pose1: the ligand’s reactive carbonyl is located 3.2 Å from S160 and is also stabilized by T-stacking with H237. However, additional interactions emerge: T88 hydrogen-bonds a nearby carbonyl oxygen, R90 and S93 donate backbone H-bonds to another carbonyl oxygen, I208 forms a π-alkyl interaction with an aromatic ring, while W159 now T-stacks both aromatic rings of the ligand. Residues N241, S238 or S92, while not directly interacting with the ligand, outline the cleft and the binding pose(s) and have positional interaction potential. In both poses, the carbonyl oxygen adjacent to the ligand’s catalytic carbon (positioned near S160) is oriented toward the backbone amide groups of Y87 and M161 in the oxyanion hole, where it is stabilized in a geometry characteristic of a transition-state stabilization. Guided by these contacts we chose as residues for mutation, from Pose 1, M161, W185, S238, W159, and from Pose 7, T88, R90, S92, S93, I208, N241, while also adding Y87 (proximity) and Q119 (groove flank).

We then used the GDEE platform to mutate the selected 12 ligand-contacting residues. In total, 14.8 million poses were generated and, to focus the analysis, we decided to prioritize the single mutant library (**Figure S1)**. Poses were filtered across four possible substrate orientations of 2HE-MHET2—its C1, C3, C4, or C6 carbonyl placed toward S160—, retaining only those that satisfied key geometric criteria relative to S160 and the oxyanion-hole residues Y87 and M161, with additional checks ensuring proper orientation and accommodation within the binding groove (**Figure S1**). Filtered poses were ranked by their predicted binding free energies and split into two symmetry groups based on the ligand positions: Poses I/II (central carbonyls C3/C4 toward S160) and Poses III/IV (outer carbonyls C1/C6 toward S160) (**Table S6**). Filtered candidates were then visually inspected to assess binding geometry and overall fit within the active site, excluding mutations known to reduce activity (e.g., S238F ^20,63,64^, and N241A ^64^). Variants with similar energies were also considered for this inspection. Ultimately, from the filtered and ranked data set, nine single mutants were selected for in vitro validation (**Table S7**).

### Screening of computationally designed variants reveals S238Y as a top performing mutation

Although FP was chosen for in silico engineering due to its reported high activity and the availability of a high resolution crystal structure ^26^, additional literature-reported PETase variants were selected to benchmark the in silico results and our experimental setup, namely the WT enzyme, TSP—which also exhibits markedly improved PET activity ^24^—and TP—the common scaffold for FP and TSP ^23^. To establish a rapid and reliable PET degradation assay, we adapted a previously reported low-volume UV absorbance method for continuous monitoring using a Nanodrop ^24^. This approach detects aromatic groups at 260 nm (BHET, MHET, TPA, and small oligomers), giving a bulk read-out of total solubilised degradation products. The extinction coefficient determined for MHET (5,464 M^−1^ cm^−1^) was used to report absorbance values as MHET equivalents (MHETEq). Experimental setup (**Figure 3A**) validation with PETase variants TS, FP, and TSP at 40 °C confirmed that absorbance trends mirrored UPLC MHET/TPA measurements, reliably capturing fold-changes and time-course activity, despite overestimating absolute concentrations (**Figure S2**). Consistent with earlier reports ^23,24,26^, TP outperformed WT and FP/TSP outperforms TP. However, our results newly demonstrate that TSP outperforms FP under all tested conditions.

**Figure 3.**
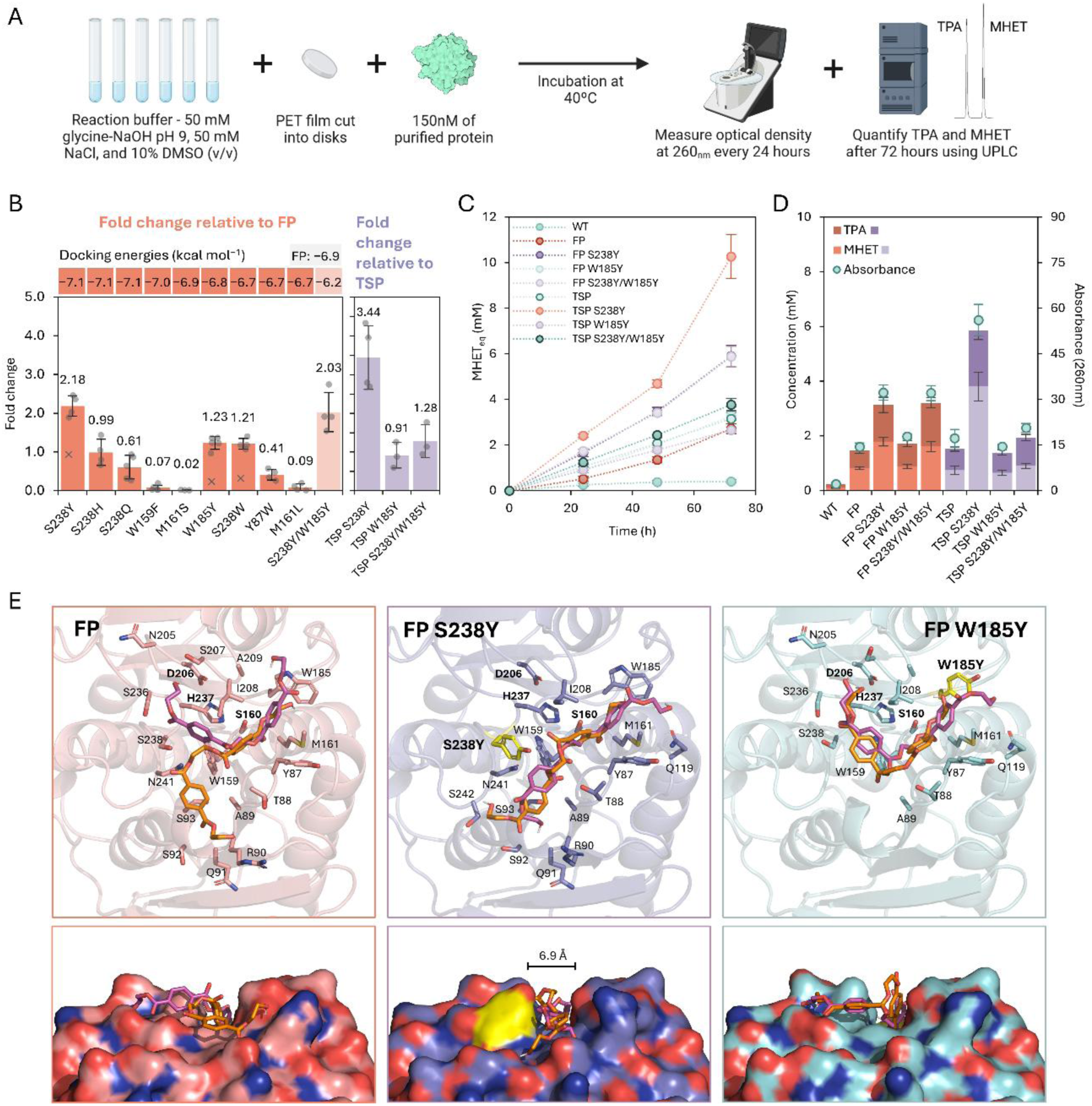
Screening of the mutant candidates selected with the GDEE workflow for enhanced enzymatic PET degradation, compared to the scaffold FastPETase (FP) and to ThermoStable-PETase (TSP), at 40 °C and 150 nM of enzyme (FP). **(A)** Schematic representation of the experimental workflow. PET film was incubated with 150 nM of purified enzyme in a glycine-NaOH buffer pH 9.0, at 40°C. Reaction progress was monitored by measuring optical density at 260 nm every 24 hours using a NanoDrop One, and monomeric products (MHET and TPA) were quantified after 72 hours using UPLC. **(B)** Mean fold change (bars) ± standard deviation (SD, n=3–6) in product formation (Abs260nm) at 72 h for each mutant relative to FP (left side, orange) and relative to TSP (right side, purple). Fold changes (mutant/scaffold) were calculated independently within each biological replicate (dots) and then averaged. Outliers are shown as crosses. Docking scores for the FP scaffold and FP mutants (top row) are shown as reference. **(C)** Time-course of soluble product accumulation over 72 h. Points represent mean Abs260nm converted into MHET equivalents (MHETEq) ± standard deviation of the mean (SEM). **(D)** Quantification of MHET and TPA by UPLC (stacked bars) after 72 h of reaction, overlaid with bulk absorbance (Abs260nm, orange circles, right y-axis). Bars represent mean concentrations ± SEM. **(E)** Cartoon (top panels) and surface lateral view (bottom panels) representations of the binding pocket of FP, FP S238Y and FP W185Y structures modelled and validated by the GDEE platform, docked with the filtered poses from the GDEE library. Visible are the ligand poses filtered by metrics rules I/II (magenta) and filtered by metric rules III/IV (orange) outputted by the GDEE pipeline.

To evaluate the newly constructed mutants, each selected FP mutant from the GDEE pipeline was assayed in vitro at 40 °C, 150 nM enzyme, and compared to its scaffold FP after 72 h (**Figure 3B, left**). Two single mutants exhibited improved activity: S238Y replacing the small, hydrophilic serine with a bulkier, hydrophobic tyrosine doubled PET degradation activity (fold change of 2.2; two-tailed t-test, *p* < 0.01), while W185Y, replacing the highly conserved and flexible tryptophan with tyrosine, delivered 1.2-fold higher soluble product (*p* > 0.05). Compared to the WT, these correspond to a fold change increase of 14.8- and 8.2-fold for FP S238Y and FP W185Y, respectively. Mutations S238H and S238W had minimal impact, whereas S238Q reduced activity by half. Mutations W159F, M161S, Y87W, and M161L showed markedly inferior activity. Time-course product accumulation over 72 h (**Figure 3C, Figure S3**) confirmed the sustained superiority of S238Y. Mutants with improved activity were selected for UPLC quantification, which validated the bulk absorbance results (**Figure 3D**). Structurally, FP (like WT PETase) retains a significantly broader active site cleft due to serine at position 238 (**Figure 3E, left**), which is a highly conserved phenylalanine in other PHE homologues (**Figure S4**). Switching to tyrosine in S238Y narrows the binding cleft and enhances its hydrophobicity (**Figure 3E, middle**). Histidine (S238H), being polar, and tryptophan (S238W), hydrophobic but lacking a hydroxyl donor, did not boost activity significantly in this position, despite also narrowing the binding cleft (**Figure S5**). Glutamine introduces a large polar side chain in S238Q that likely disrupts substrate packing, leading to the observed activity loss (**Figure S5**). Mutation W185Y left the shape of the catalytic grove largely unchanged (**Figure 3E, right**).

Combining mutations S238Y and W185Y yielded a double mutant scoring −6.2 kcal mol^−1^ in GDEE and that resulted in no further benefit over S238Y (**Figure 3D**), maintaining an activity similar to FP S238Y (*p* < 0.01). Furthermore, to test whether the benefits of the identified beneficial mutations extended to the more active TSP scaffold, we introduced S238Y and W185Y into TSP (**Figure 3B**, **right**), which had already outperformed FP (**Figure S2**). In TSP, S238Y tripled activity (3.4-fold, *p* < 0.01), W185Y was neutral (0.9-fold) and S238Y/W185Y had a modest increase in activity, (**Figure 3A**). Time-course and UPLC analyses confirmed TSP S238Y as the strongest performer (**Figure 3B,C**).

Moreover, to assess performance under lower pH, the best FP- and TSP-derived variants were assayed at pH 8.0 and 7.5 (**Figure S6**). As expected for the enzyme PETase—which has optimal performance around pH 9.0, with a steep decrease at lower pH ^15,64^—activity declined with decreasing pH: FP and FP S238Y lost most activity at pH 8.0, whereas FP S238Y/W185Y retained 57 % of its pH 9.0 activity. TSP S238Y retained 54 % at pH 8.0 and 33% at pH 7.5. In summary, at 40 °C FP S238Y showed a 14.8-fold increase in activity relative to PETase WT based on absorbance, corresponding to 9.4-fold more TPA and 20-fold more MHET by UPLC. TSP S238Y reached a 25.8-fold increase measured by absorbance compared to the WT (14.4-fold more TPA and 42.6-fold more MHET) and retained an appreciable level of activity at lower pH.

### Kinetic evaluation shows that S238Y reduces concentration inhibition

While bulk absorbance and endpoint quantification are useful for evaluating PETase activity, they offer limited insight about how cavity mutations reshape catalysis. However, modelling with conventional Michaelis-Menten (MM) kinetics is not suitable in this case, as the model assumes excess of soluble substrate, whereas PET hydrolysis occurs at the solid-liquid interface where substrate accessibility changes over time ^40,65^. As the PET surface is degraded, velocity depends on how many enzyme molecules adsorb and react productively. Langmuir adsorption-based and inverse MM models have been proposed to better capture this interfacial behaviour ^40,65–67^ by assuming that reaction velocity increases with enzyme concentration until saturation is reached, reflecting the proportion of enzyme molecules productively bound at accessible sites on the polymer surface.

To investigate how the S238Y and W185Y mutations individually influence the activity and kinetics of FP, and how S238Y affects those of TSP, we measured initial reaction rates from 25 to 400 nM enzyme using a PET load of 8.23 g L^−1^ (0.612 cm^2^ surface area), corresponding to 40.8 to 653.6 nM cm^−2^ enzyme loading. Consistent with previous work ^24,65,67^, initial reaction rates increased up to an optimum around 100–150 nM enzyme—about 100 nM for the FP mutants and 100–150 nM for TSP mutants—and then mostly declined due to enzyme concentration inhibition (**Figure 4A,B; Figure S7**). This decline was particularly evident for FP at 300 and 400 nM and FP W185Y at 400 nM: there was virtually no activity during the first 4 hours, preventing the calculation of an initial rate. At the optimal enzyme concentration, FP S238Y and FP W185Y were 1.7-and 1.5-times faster than FP; beyond 150 nM both mutants resisted inhibition better, having ∼4-fold higher activity at 200 nM. FP S238Y alone retained measurable turnover at 400 nM (**Figure 4A**). TSP S238Y showed a broader optimal range (100–200 nM), reduced concentration-dependent inhibition (65.2 ± 14.7 % for TSP, compared to 10.3 ± 9.2 % for TSP S238Y), and a higher catalytic rate than TSP—2-fold at 150 nM and 5.5-fold at 400 nM. When compared to the TSP mutants, FP and its mutants showed greater inhibition, from 83.9 % to 88 %. Since initial rates do not always reflect long-term performance, PET hydrolysis was monitored over the next 72 hours (**Figure 4C,D**). The initial inhibition at high enzyme concentrations diminished over time for FP S238Y and FP W185Y (**Figure 4C**) and was largely alleviated in TSP, whose inhibition dropped from 69.8 % at 4 h to 11.1 % at 72 h. For FP W185Y, activity and product accumulation remained lower than that of FP S238Y. TSP S238Y maintained low inhibition levels throughout (10.6 % at 4 h and 15.8 % at 72 h) and showed an exponential increase in product accumulation across 150–400 nM. At 200 nM, TSP S238Y’s product yield was 22.6-fold higher than TSP (**Figure 4D**).

**Figure 4.**
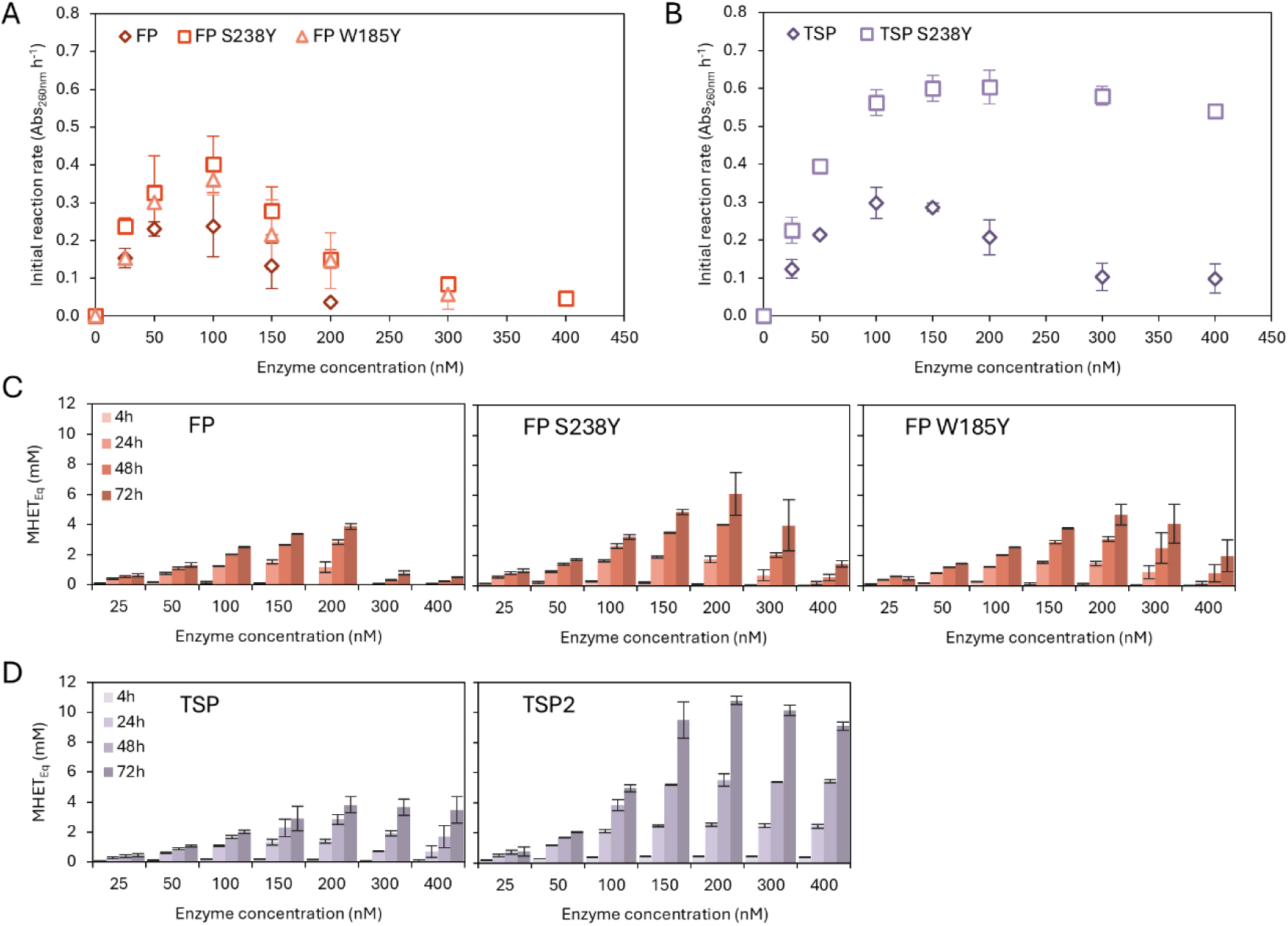
Kinetic profiles and time-course analysis of PET film degradation by FP and TSP backgrounds and corresponding engineered best performing single variants. **(A)** Initial reaction rates (product formation in terms of absorbance at 260 nm per hour) over 4 hours for different enzyme concentrations (25, 50, 100, 150, 200, 300 and 400 nM) for FP and FP-derived single mutants (S238Y, W185Y), at 40 °C. **(B)** Initial reaction rates for TSP and TSP-derived mutant TSP2 (TSP S238Y) under the same conditions as in (A). **(C)** Absorbance-based MHETEq quantification of soluble PET degradation products produced by different enzyme concentrations, for the FP background and its mutants over 72h. **(D)** Absorbance-based MHETEq quantification of soluble PET degradation products produced by different enzyme concentrations, for the TSP and TSP S238Y over 72h. Bars and data points represent mean ± standard deviation (SD) from biological duplicates (n = 2).

Since activity decreased at higher enzyme loadings, most variants could not be fitted to the inverse MM model (**Equation 1**). TSP S238Y, being least affected by inhibition, was fitted up to the start of activity decline (200 nM), yielding an ^inv^KM = 53.1 ± 10.7 nM and an ^inv^Vmax = 0.8 ± 0.1 Abs260nm h^−1^. Here, ^inv^Vmax reflects the point at which the PET surface becomes saturated with enzyme, mechanistically indicating the total number of accessible binding sites on the solid substrate, while ^inv^KM reflects the enzyme concentration required to half-saturate the surface and is a proxy for affinity for the PET surface ^66^.

### Molecular dynamics simulations reveal stabilizing effects

To simulate the behaviour of the tested variants and scaffold FP with the ligand model, we performed MD simulations starting from the modelled GDEE variant and the corresponding docking pose ligand. RMSD of Cα atoms revealed an overall stable structure for FP (0.78 ± 0.02 Å) with only a few variants deviating from this stability. In particular, mutation M161S (fold change = 0.09) displayed the highest RMSD (1.06 ± 0.16; **Figure S8**). Throughout the simulations, the ligand generally remained in the binding cleft, only occasionally dissociating (data not shown). RMSD analysis of cleft residues (∼4 Å around the ligand) revealed higher values in the worst in vitro-performing mutants, suggesting that increased local flexibility and deformation of the active site correlate with reduced activity (**Figures S9 and S10**).

To identify key ligand-interacting residues, we filtered MD frames where at least one of the ligand’s catalytic carbons is positioned within 6 Å of the catalytic serine and identified those residues in close contact (≤ 4 Å) with the ligand. This revealed recurrent contact hotspots such as loop segments G86–A89 and W159–W161, located adjacent and bellow the binding cleft, as well as residues W185, I208 and H237 (**Figure 5A**). Contact frequencies for several residues correlated positively with in vitro activity, most strongly in position 238 (*r* = 0.80; *p* = 0.003; **Figure 5B**), followed by T88 (*r* = 0.70; *p* = 0.01), A89 (*r* = 0.64; *p*=0.033) and W159 (*p* = 0.62; *r* = 0.042). This observation aligns with GDEE filtered docking poses, where S238Y creates a protrusion or wall that may improve PET chain accommodation. In a representative MD snapshot of scaffold FP (Error! Reference source not found.**C)**, the binding cleft is wider and the ligand bends more loosely along the cleft plane – a configuration also observed among mutants M161L, M161S, W159F, W185Y and Y87W, but not in S238Y or S238Y/W185Y. In these latter mutants, tyrosine at position 238 forms a barrier that narrows the cleft, markedly increasing ligand contact with Y238; a similar pattern occurs in S238W. Mutants S238H and S238Q also introduce bulkier side chains, but neither produces such a substantial barrier. Additionally, per-donor hydrogen bond occupancy analysis between ligand acceptors and the cleft donors revealed an increased H-bond formation with mutated residues at position 238 (**Figure S11**). Consistent with its central catalytic role, S160 had the highest ligand H-bond occupancy across variants, except in M161S, which correlates with its abolished in vitro activity.

**Figure 5.**
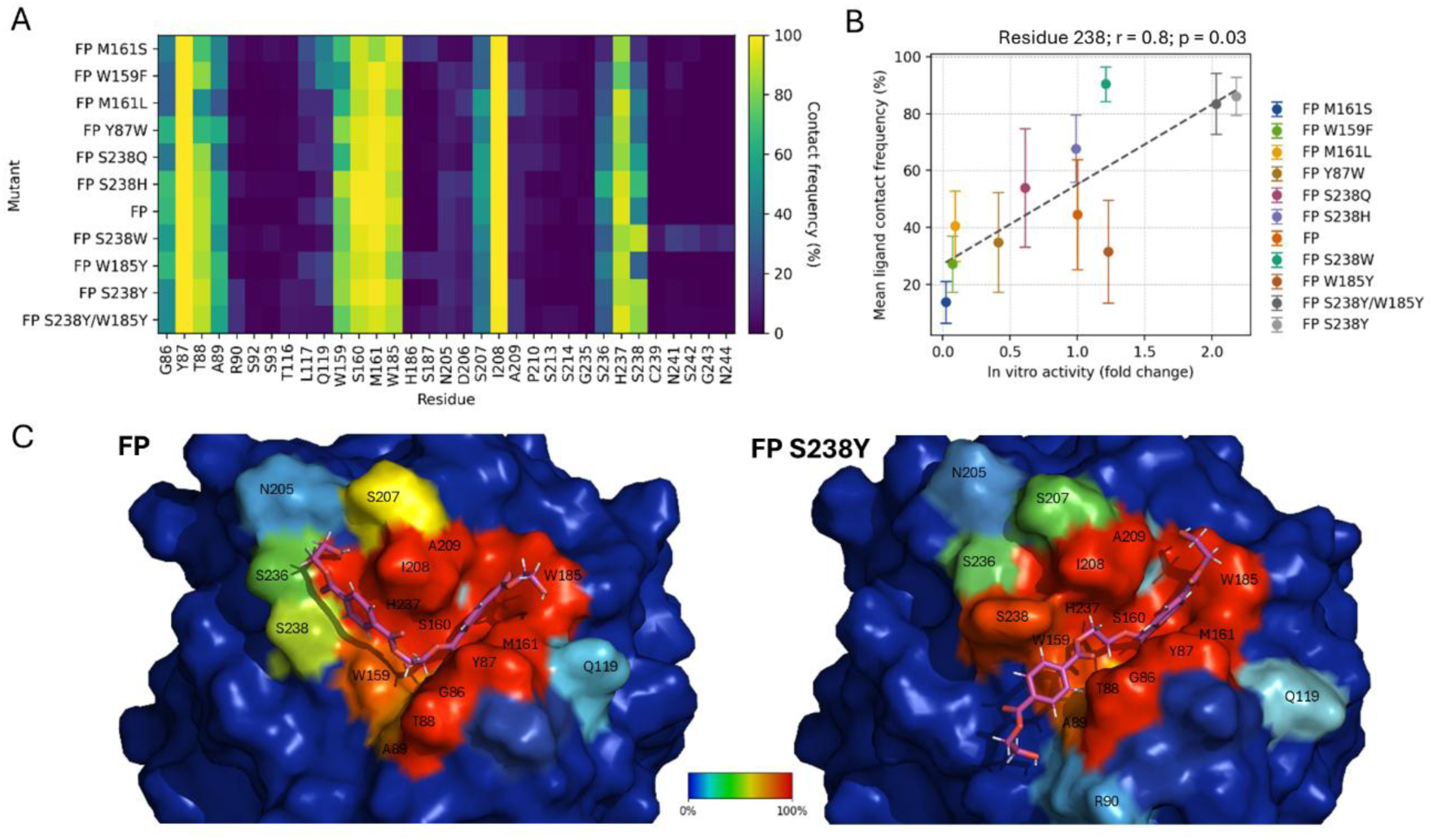
Analysis of the Molecular Dynamics (MD) simulations performed for the FP variants (0.2 µs; n=5). **(A)** Heatmap of ligand contact frequencies for active-site residues across PETase mutants. The y-axis lists each mutant, and the x-axis lists non-muted residue identities and positions. Colours indicate the mean percentage across runs of the number of MD simulation frames in which the shortest heavy-atom distance between the residue and the ligand was ≤ 4 Å. Only frames where the catalytic S160-ligand distance was ≤ 6 Å were considered for percentage calculation. Residues from the active site cleft selection with negligible contact frequencies are not shown. **(B)** Relationship between in vitro activity and mean residue 238-ligand interaction for each variant with error bars representing 95% confidence intervals (± 2σ) from 1000 bootstrap resampling iterations. Dashed lines indicate least-squares linear fits (Backbone fit: y = 28.0x + 27.1; *r*=0.8; *p*=0.03). **(C)** Representative MD simulation structures (protein and ligand) of FP and FP S238Y with residue-specific contact frequencies encoded in the B-factor field and visualized as a colour gradient. The colour bar indicates the colour from lower to higher contact frequency percentage (0–100%).

Furthermore, we analysed π-stacking interactions between the ligand’s phenyl rings and the aromatic rings of S238Y, S238W, and S238H mutants. The ligand formed parallel-displaced interactions with tyrosine and tryptophan, indicating the establishment of a new interaction type that may help stabilize the ligand, whereas S238H displays sporadic T-shaped interactions (**Figure S12A and B**). The π–stacking analysis of W185/Y185 with the ligand revealed predominantly T-shaped geometries, with their frequency increasing in W185Y and S238Y/W185Y (**Figure S12C and D**). Finally, we analysed the χ^1^ and χ^2^ dihedral angles of tryptophan and tyrosine in position 185 and found that the conformer B predominated both in the presence and absence of the ligand (data not shown). In W185Y, the mean χ^2^ angle decreased slightly 88.5° ± 2.0° compared with 96.2° ± 1.5° for FP; however, visual inspection confirmed that both values correspond to highly similar orientations, consistent with conformer B.

## DISCUSSION

The aim of this work was to identify PETase mutants with improved PET hydrolytic activity at 40 °C by exclusively targeting residues at the active site cavity. FP was selected as the engineering scaffold for its enhanced thermostability and activity in this temperature range—conferred by distal mutations without altering the catalytic site ^26^—compared to other known PHEs ^68^ such as DuraPETase ^25^ and LCC^ICCM^ ^18^. We selected a flexible 2HE-MHET2 docking ligand, since longer segments such as 2HE-MHET4^17,20^ or 2HE-MHET3 have given unreliable and unstable poses.

Using the GDEE platform ^29^, we identified several potentially beneficial mutations, tested them in vitro, and found S238Y as a key beneficial mutation in the FP background, yielding a mutant with double the activity of FP on PET disks and higher initial reaction rate. Addition of W185Y—which substituted the highly conserved tryptophan (**Figure S4**) by tyrosine, yielding a 1.2-fold activity increase on the FP background—to FP S238Y produced negligible additional gain. Introducing S238Y into the higher-performing TSP background yielded TSP S238Y, which outperformed all variants under every condition, and showed 3.4-fold activity increase over TSP. TSP S238Y maintained high performance over a range of enzyme concentrations and showed reduced concentration-dependent inhibition, a behaviour that is otherwise a known limitation of mesophilic PETases ^15,24,40,65,67^. This inhibition is likely caused by enzyme crowding (i.e., inhibitory enzyme-enzyme interactions in the PET surface), leading to reduced surface accessibility and increased enzyme rigidity ^65,69^. This effect may be attenuated in TSP S238Y due to faster PET surface turnover that limits surface saturation. Further addition of mutation W185Y to TSP S238Y yielded a double mutant with only 1.3-fold activity increase compared to TSP alone, implying that W185Y is deleterious in this background. When compared to PETase WT, FP S238Y showed a 14.8-fold increase in activity based on absorbance, corresponding to 9.4-fold more TPA and 20-fold more MHET by UPLC, while TSP S238Y reached a 25.8-fold increase by absorbance (14.4-fold more TPA and 42.6-fold more MHET).

We also examined pH effects on activity, as PETase is known to exhibit optimal activity under alkaline conditions ^15,64^. At physiological pH, PETase’s high 9.6 isoelectric point and strong dipole ^22^ give it a positive charge, likely promoting electrostatic interactions with negatively charged PET surfaces that may hinder product release and turnover. At pH 9, improved charge balance and optimal protonation of catalytic residues—such as the catalytic H237 which has predicted p*K*a of 6.78 (PDB: 7SH6; PropKa 2.0)—may enhance catalysis. While TSP S238Y and FP S238Y/W185Y followed this trend, both still retained considerable activity at pH 7.5–8.0, indicating enhanced pH tolerance. This may result from mutation-induced changes in the active-site electrostatics that reduce sensitivity to pH-dependent charge shifts, potentially stabilising catalytically productive conformations.

Other GDEE mutations gave poor outcomes, for example, W159F nearly abolished activity, and mutations in the oxyanion hole residues were similarly detrimental: replacing the highly conserved M161 (**Figure S4**) with serine or leucine eliminated activity, and Y87W reduced turnover to 40 % of FP. Previous reports of substitutions in the oxyanion hole had similar outcomes: Y87F led to reduced PET bottles degradation ^70^ even though phenylalanine has a degree of conservation in this position (**Figure S4**), while M161Q abolished activity ^71^.

MD simulations and subsequent RMSD analysis showed that FP and its variants were overall structurally stable, with only a few variants deviating from this trend. However, residues at the active site cleft displayed a correlation between increased loop instability, reflected by higher local RMSD values, and reduced in vitro activity. This suggests that mutations in these low-performing variants introduced elevated local flexibility and subtle deformation of the catalytic groove, which in turn correlated with decreased turnover. These findings support a model in which higher mobility within the cleft compromises catalytic efficiency.

A detailed analysis of ligand–residue contacts frequencies in the binding cleft during MD simulations revealed a strong positive correlation with in vitro activity for several residues, particularly at position 238. Variants exhibiting higher contact frequency at this position consistently showed enhanced catalytic rates. Introduction of aromatic side chains at position 238 (S238Y, S238W) promoted new parallel-displaced π–π stacking interactions with the ligand’s phenyl rings. These aromatic interactions were accompanied by transient hydrogen bonds between the ligand and the mutated residues at position 238, further stabilizing the complex. In contrast, S238H formed weaker, sporadic T-shaped interactions that did not translate into significant activity gains. Simulations also revealed that the bulker tyrosine in S238Y introduces a steric barrier that narrows the binding groove and interacts directly with the ligand, potentially lowering ligand escape rates. Altogether, these findings support a stabilisation model in which aromatic and hydrophobic interactions with the ligand at position 238—particularly those mediated by tyrosine—play a central role in promoting ligand positioning and catalytic efficiency. Finally, the modest activity increase observed for W185Y likely reflects subtle changes in conformational flexibility while retaining the functional geometry of the native tryptophan. MD simulations confirmed that tyrosine at position 185 preserves the B conformer and continues to supports substrate interaction, while also modestly increasing the frequency of π–π contact with the ligand.

Most positions mutated in this work are conserved across PHEs, except S238, which is a serine in PETase but a highly conserved phenylalanine in most homologues (**Figure S4**) —for example, F255 in Cut190 ^72^, reported as important for hydrophobic PET binding, and F243 in LCC or F209 in TfCut2, both predicted to interact with a docked 2PET structure while S238 in PETase does not ^21^. Serine in this position contributes to PETase’s unusually wide catalytic cleft —about three times wider than that of TfCut2 ^22^ —thought to facilitate bulky PET binding ^64^. Attempts to restore phenylalanine generally reduced activity ^20,63,64^, with only modest gains when combined with W159H ^22^, and are comparable to the limited improvement of S238W observed here relative to FP. In contrast, S238Y may provide a unique advantage by narrowing and increasing the hydrophobicity of the cleft while enabling hydrogen bonding. Only few PHEs carry a tyrosine at the position equivalent to PETase’s S238 (e.g., Ple628 and PE-H ^73,74^, PsCut and PbauzCut ^65^), but most have different cleft architectures due to loop variations that narrow the groove. In PE-H, mutating Y250 (S238-equivalent) to serine widened the cleft and boosted activity ^74^, though structural differences limit direct comparison with PETase. Together, these comparisons and observations highlight residue 238 as a key modulator of cleft geometry and PET affinity, underscoring the value of resolving the TSP S238Y structure to elucidate the structural basis of its enhanced activity.

While most engineering efforts have targeted thermostability, fewer have focused on modifying the catalytic site to directly enhance degradation efficiency. Exceptions include the study by Son et al., where structural bioinformatics was used to compare active site residues across four PETase homologues, identifying S242T and distal N246D as beneficial mutations that improved activity by 1.5-fold and 2.4-fold, respectively, at 37 °C compared to the WT ^70^; and Sevilla et al., where active-site engineering guided by multiple sequence alignment of the WT PETase with 39 PETase homologues led to the selection of three mutations: I208V, S238Y, and distal N212A ^75^. In the latter study, published during our experimental validation, PETase S238Y was reported to exhibit a 20 % increase in binding affinity in MD simulations, using the same substrate analogue employed here. Enhanced PET surface modification was observed on post-consumer bottles samples, supported by SEM, AFM, and DSC analyses at 30 °C after 72 h, compared to WT. However, no enzymatic in vitro depolymerization or kinetic results were presented ^75^. Our study also identified S238Y, but through a fully automated, high-throughput, structure-guided pipeline. This parallel finding reinforces the mutation’s relevance, as it resulted from an analysis of a great variety of mutants, and highlights the robustness of our in silico approach. By inserting this mutation of the powerful FP scaffold and additionally quantifying degradation products and initial reaction rates, we provided a thorough functional evaluation, both validating S238Y’s potential under conditions relevant to industrial PET recycling and developing a highly active PETase mutant enzyme.

Finally, a few considerations are worth making. First, tests were performed on amorphous PET in disk form, which restricts hydrolysis to surface layers. Increasing surface area via milling or grinding, and reducing crystallinity through amorphization, enhance degradation ^17,40,76^, and will likely be essential for industrial-scale efficiency ^11^, as even highly active PHEs cannot efficiently degrade high-crystallinity PET fragments ^26^. Overall, post-consumer PET waste treatment still presents many challenges, and critical industrial parameters must be considered ^68^, as it often consists of mixed, heterogeneous fractions with varying crystallinity and additives that can hinder enzymatic processing and increase costs ^11^. Secondly, PETase has limited ability to degrade MHET ^15,76^, resulting in its accumulation as a primary hydrolysis product, where it can act as an inhibitor of PETase activity ^40^. This underscores the future benefit of applying a dual-enzyme system combining PETase with MHETase to achieve complete MHET depolymerization. Among the final monomers, TPA can be recovered at high purity ^18,26^, while EG is suitable for microbial assimilation and upcycling into value-added compounds via metabolic engineering ^77^. Together, these factors highlight the opportunities and the current limitations of enzymatic PET recycling. Future efforts will need to integrate enzyme engineering with process optimization to address substrate heterogeneity, surface accessibility, and MHET accumulation. Ultimately, combining highly active PETase with improved versions of MHETase, alongside PET pretreatment strategies, will be key to advancing industrial-scale PET bioprocessing.

## CONCLUSION

In this work we employed a structure-guided, automated protein engineering platform, called GDEE, to generate, in a high-throughput manner, a large library of PETase mutant candidates targeting residues at the active site, docked with a PET substrate analogue. This was followed by a rigorous filtering and selection process, enabling the identification of a smaller set of promising mutations for in vitro validation. By streamlining the experimental screening of engineered PETase variants, this approach accelerates the discovery of enzymes with potential applications in plastic biodegradation and recycling, while significantly reducing laboratory workload. We identify S238Y as a key mutation that enhances PET film degrading performance at 40 °C when inserted in two of the most active PETase variants reported to date: 2.2-fold increase in the FP scaffold and 3.4-increase in the TSP background. FP S238Y showed a 14.8-fold increase in activity based on absorbance compared to PETase WT, translating into 9.4-fold more TPA and 20-fold more MHET by UPLC, while TSP S238Y reached a 25.8-fold increase (14.4-fold more TPA and 42.6-fold more MHET). This mutation not only improved long-term product accumulation but also enhanced catalytic efficiency and resistance to enzyme concentration inhibition, especially in the TSP scaffold. The engineered TSP S238Y variant specially stands out as a highly promising candidate foundation for further rational design and for further development in mesophilic, scalable and efficient PET degradation systems.

## AUTHOR INFORMATION

### Corresponding Author

Sofia Ferreira – Instituto de Tecnologia Química e Biológica António Xavier, Oeiras, Portugal;

Cláudio M. Soares – Instituto de Tecnologia Química e Biológica António Xavier, Oeiras, Portugal;

Isabel Rocha – Instituto de Tecnologia Química e Biológica António Xavier, Oeiras, Portugal;

### Present Addresses

Alexandra Balola - Instituto de Tecnologia Química e Biológica António Xavier, Oeiras, Portugal

Sofia Ferreira - Instituto de Tecnologia Química e Biológica António Xavier, Oeiras, Portugal

Caio Silva Souza

Diana Lousa - Instituto de Tecnologia Química e Biológica António Xavier, Oeiras, Portugal

João Correia - Instituto de Tecnologia Química e Biológica António Xavier, Oeiras, Portugal

Cláudio M. Soares - Instituto de Tecnologia Química e Biológica António Xavier, Oeiras, Portugal

Isabel Rocha - Instituto de Tecnologia Química e Biológica António Xavier, Oeiras, Portugal

### Author Contributions

**AB** performed all experiments, analysed the data, and wrote the manuscript. **CMS**, **CSS**, and **JC** developed the computational engineering platform GDEE. **CMS**, **CSS**, and **DL** provided support with the setup and execution of the in-silico engineering rounds. **SF**, **CMS** and **IR** supervised all experiments and reviewed and edited the manuscript.

### Notes

The authors declare no competing financial interest.

## ACKNOWLEDGMENTS

We would like to thank the Protein Purification & Characterization and the Protein Purification & Characterization Research facilities from ITQB.

## ABBREVIATIONS

PET – Polyethylene Terephthalate, TP – ThermoPETase, FP – FastPETase, TSP – ThermoStablePETase, EG – Ethylene Glycol, TPA – Terephthalic Acid

