## Supplementary Information for "Computer-guided enzyme engineering of PET hydrolase mutants towards improved PET affinity"

### Supporting Figures

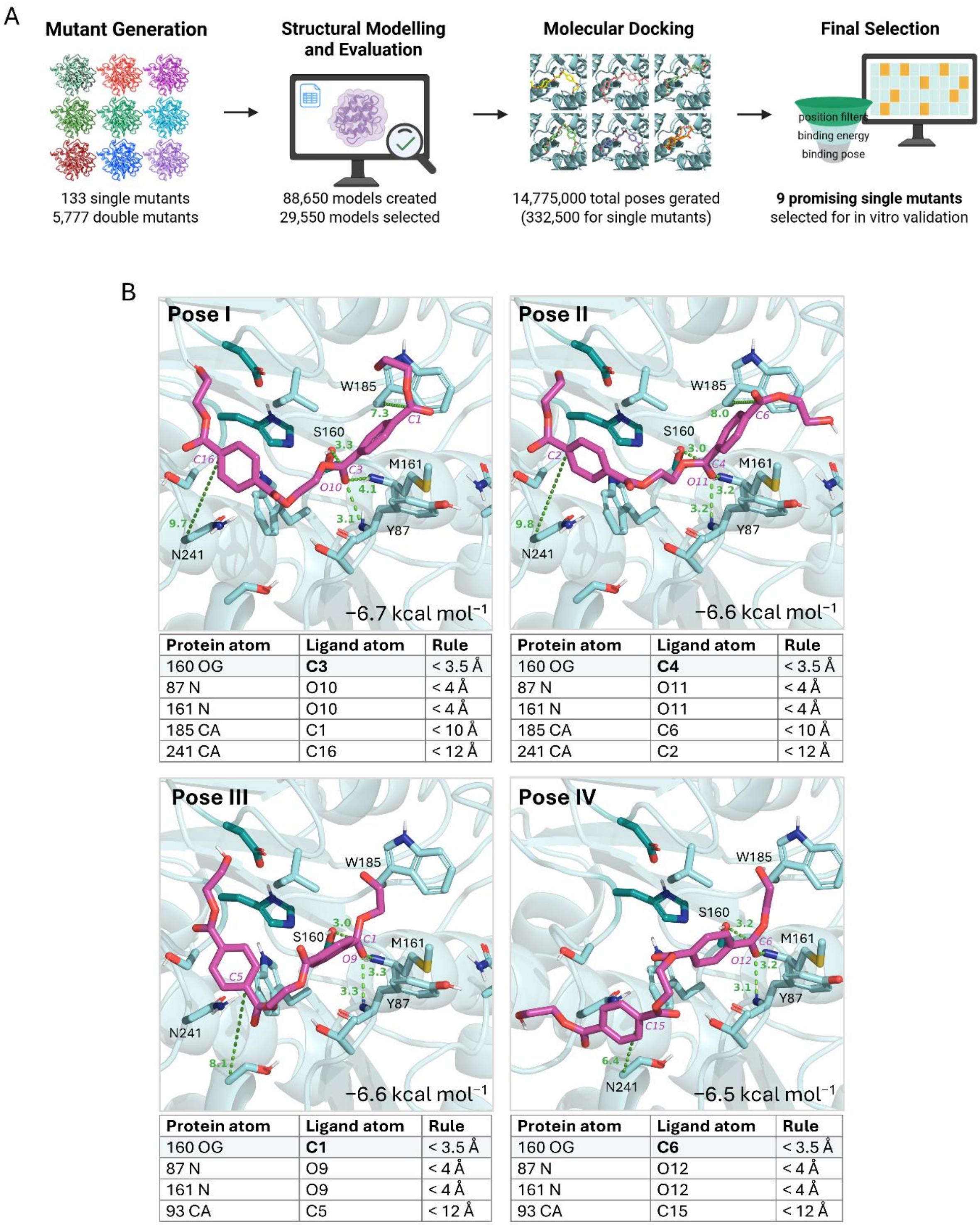

Pose I

| Protein atom | Ligand atom | Rule |
|---|---|---|
| 160 OG | **C3** | < 3.5 Å |
| 87 N | O10 | < 4 Å |
| 161 N | O10 | < 4 Å |
| 185 CA | C1 | < 10 Å |
| 241 CA | C16 | < 12 Å |

Pose II

| Protein atom | Ligand atom | Rule |
|---|---|---|
| 160 OG | **C4** | < 3.5 Å |
| 87 N | O11 | < 4 Å |
| 161 N | O11 | < 4 Å |
| 185 CA | C6 | < 10 Å |
| 241 CA | C2 | < 12 Å |

Pose III

| Protein atom | Ligand atom | Rule |
|---|---|---|
| 160 OG | **C1** | < 3.5 Å |
| 87 N | O9 | < 4 Å |
| 161 N | O9 | < 4 Å |
| 93 CA | C5 | < 12 Å |

Pose IV

| Protein atom | Ligand atom | Rule |
|---|---|---|
| 160 OG | **C6** | < 3.5 Å |
| 87 N | O12 | < 4 Å |
| 161 N | O12 | < 4 Å |
| 93 CA | C15 | < 12 Å |

**Figure S1.** Overview of the GDEE high-throughput protein engineering workflow and of the metrics selected to filter the results library. (A) Schematic representation of the GDEE high-throughput protein engineering workflow used for automated mutant generation, structural modelling, docking and library filtering of thousands of protein variants with the docked ligand.

Library sizes obtained in this study are indicated. Scheme created with BioRender.com. (B) Representative docking poses of the ligand 2HE-$MHET_2$ bound to the active site of FastPETase (FP; PDB ID: 7SH6) highlighting the key metrics/distances used to filter the GDEE results library. Poses I–IV illustrate one of the four main productive binding poses, where one of the four catalytic carbonyl carbons of the ligand aligns with the catalytic S160, identified from docking results. The ligand is shown in pink sticks; selected distances (shown in green, in angstroms [Å]) reflect the filtering criteria. These metrics and the corresponding threshold rules are summarized in the accompanying table.

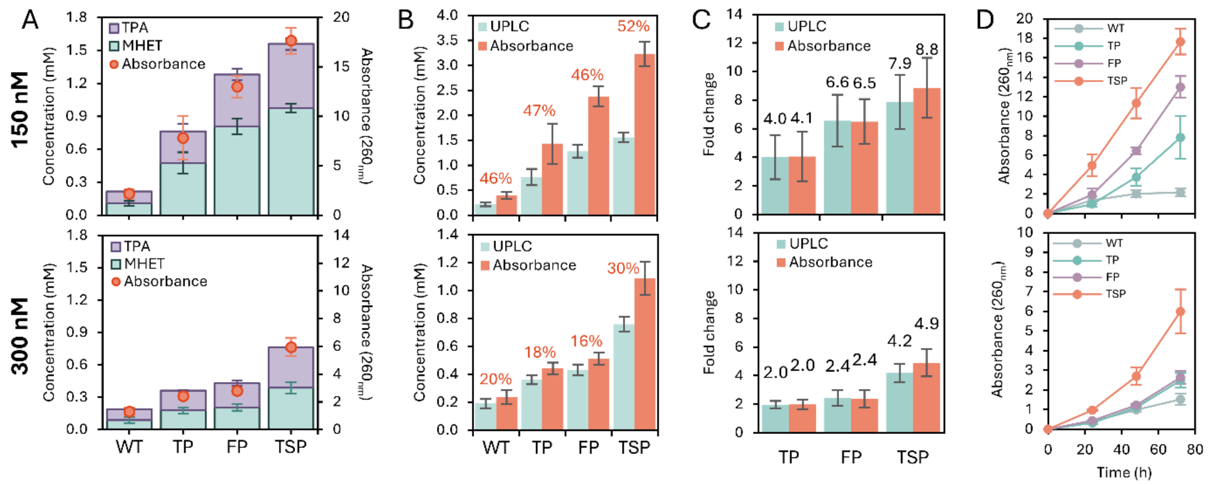

**Figure S2.** Comparison of UPLC and absorbance-based quantification of PET hydrolysis products for PETase wild type (WT), ThermoPETase (TP), FastPETase (FP) and Thermostable PETase (TSP) at 40 °C and two different enzyme concentration conditions. (A) Concentration of TPA and MHET (stacked bars) measured by UPLC, overlaid with bulk absorbance measurements at 260 nm (orange circles) for reactions conducted with 150 nM (top) after 72 hours and 300 nM (bottom) of enzyme. (B) Comparison of total soluble products quantified by UPLC (green bars) and absorbance at 260 nm at 72 hours, converted to MHET equivalents (MHETEq; orange bar), for reactions with 150 nM (top) and 300 nM (bottom) enzyme concentrations. Overestimated amount of products by the absorbance method is indicated as percentages. (C) Fold-change in total product formation relative to the WT, as quantified by UPLC (green bars) and bulk absorbance converted to $MHET_{Eq}$ (orange bars), for 150 nM (top) and 300 nM (bottom) enzyme concentrations. (D) Time course of absorbance at 260 nm over 72 hours for WT and selected PETase variants at 150 nM (top) and 300 nM (bottom) enzyme concentrations. Data are presented as the mean across experiments ± standard deviation of the mean (SEM) (n=3).

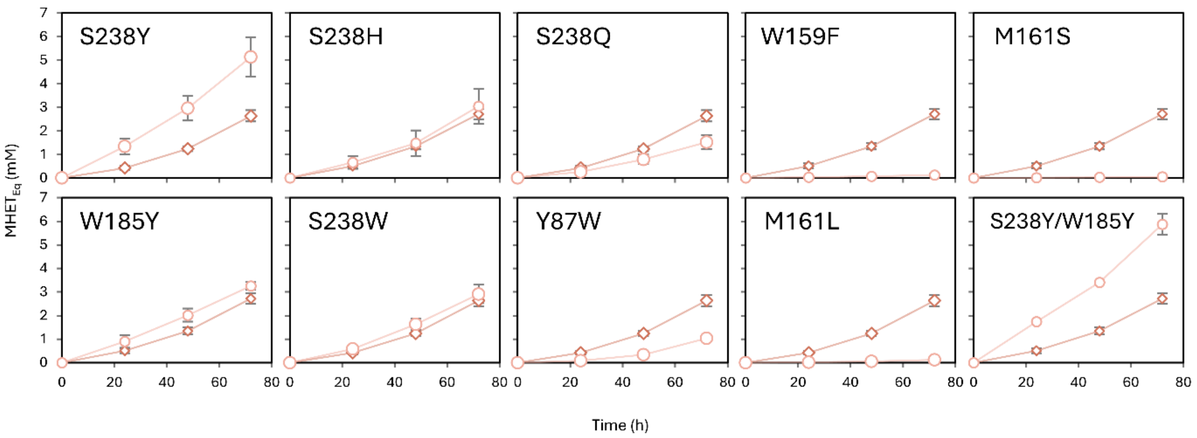

**Figure S3**. Time-course of soluble product accumulation over 72 h for FastPETase (FP; diamons), and mutant enzymes (circles). FP is shown in all graphics as reference. Points represent mean absorbance at 260 nm converted into MHET equivalents ($MHET_{Eq}$) ± SEM across biological replicates. Each enzymatic mutant was tested with 4–6 biological replicates (n=4 for Y87W, W159F, M161L, S238H, S238Y/W185Y; n=5 for FP, S238Y, S238W, S238; and n=6 for FP). Differences in replicate number were due to experimental availability, but all replicates were treated identically.

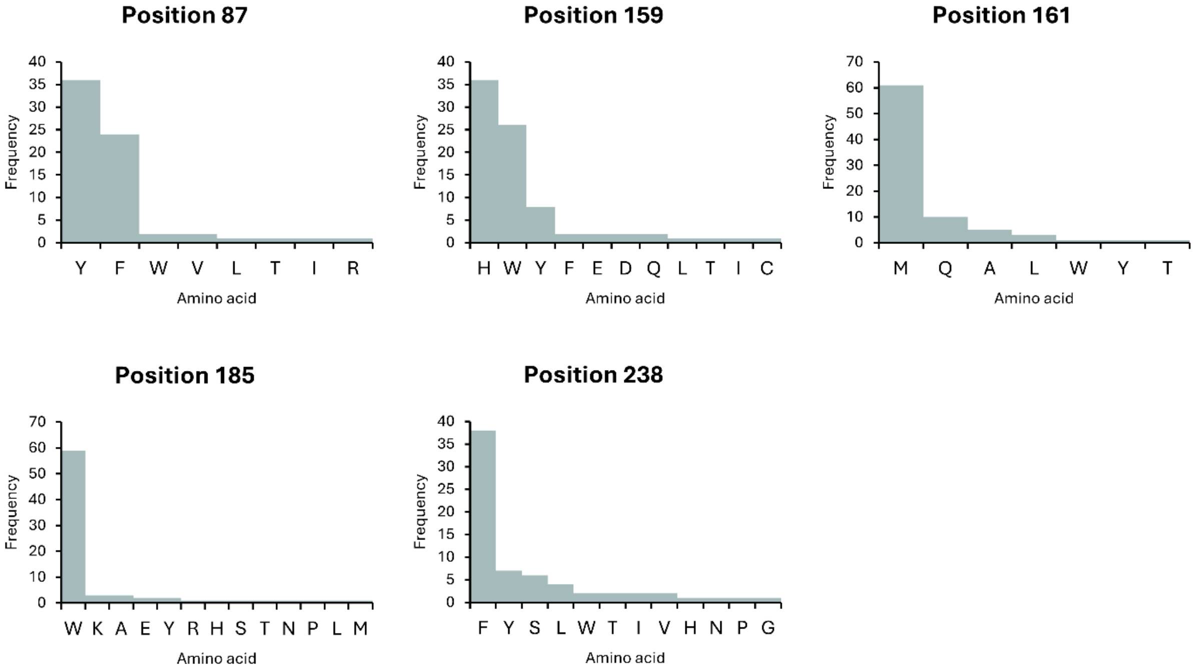

**Figure S4.** Distribution of amino acid residue frequencies at selected positions in the PETase homologues multiple sequence alignment. Each panel shows the frequency of amino acid residues observed at a specific alignment position across homologous PET hydrolase sequences with more than 45 % sequence identity.

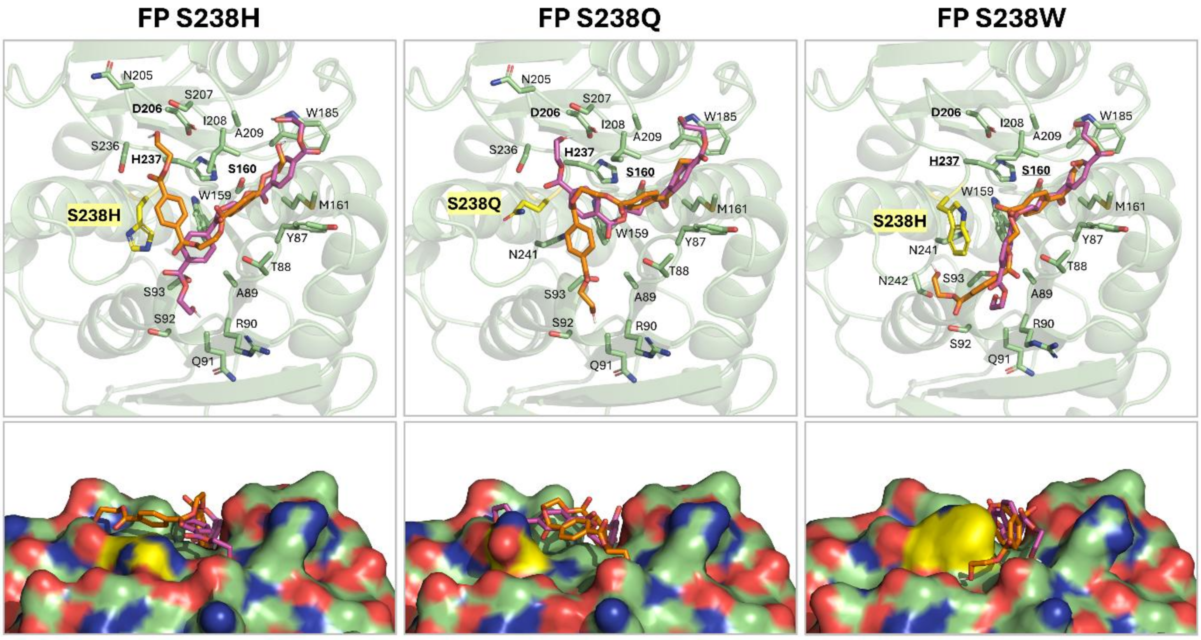

**Figure S5**. Cartoon (top panels) and surface lateral view (bottom panels) representations of the binding pocket of FP S238YW and FP W185Q structures modelled and validated by the GDEE platform, docked with the filtered poses from the GDEE library. Visible are the ligand poses filtered by metrics rules I/II (magenta) and filtered by metric rules III/IV (orange) outputted by the GDEE pipeline.

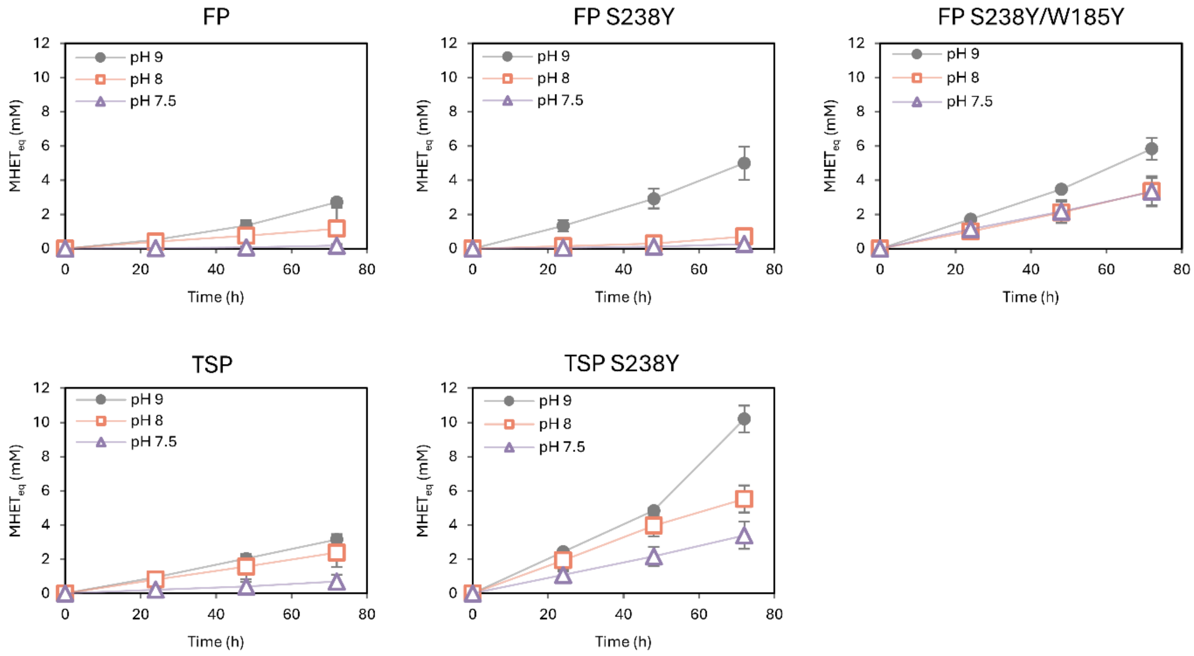

**Figure S6**. Comparison of bulk-quantified soluble product concentrations at 72h, converted to MHET$_{Eq}$, across two reaction pH conditions (pH 8.0 and pH 7.5). Points represent mean concentrations across two biological replicates ± standard deviation (SD; n = 2). For reference, values obtained previously at pH 9.0 (grey circles) are shown ± SEM (n = 4-6).

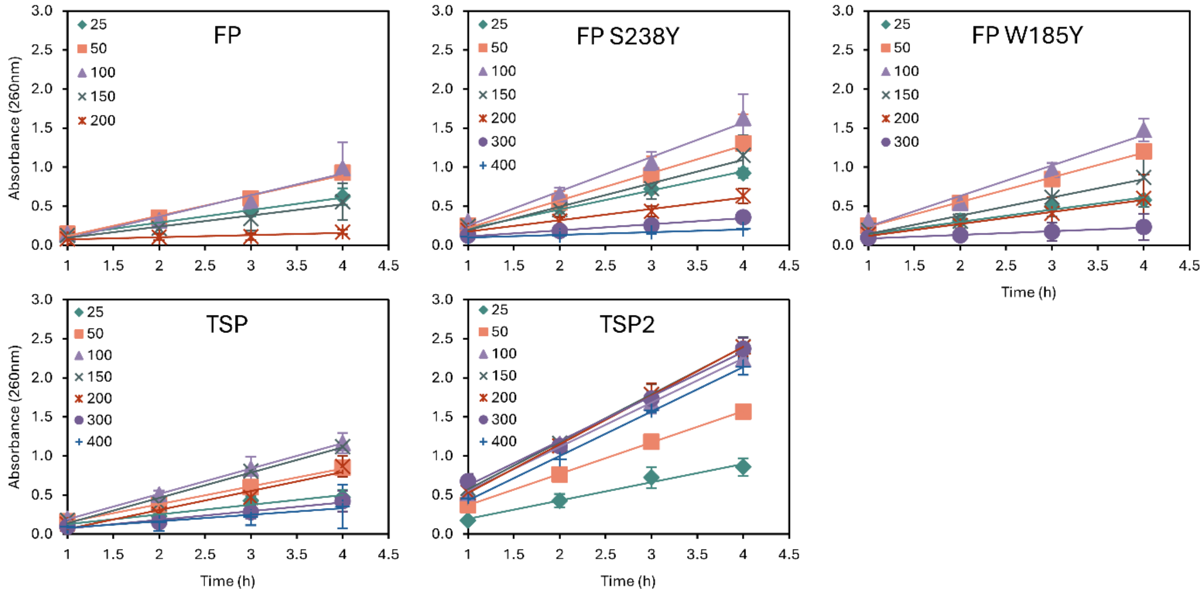

**Figure S7**. Initial reaction rates measured over 4 h in terms of bulk absorbance at 260 nm, of PET film degradation by FP and TSP backgrounds and corresponding engineered best performing single variants. Points represent mean ± standard deviation (SD) from biological duplicates (n = 2).

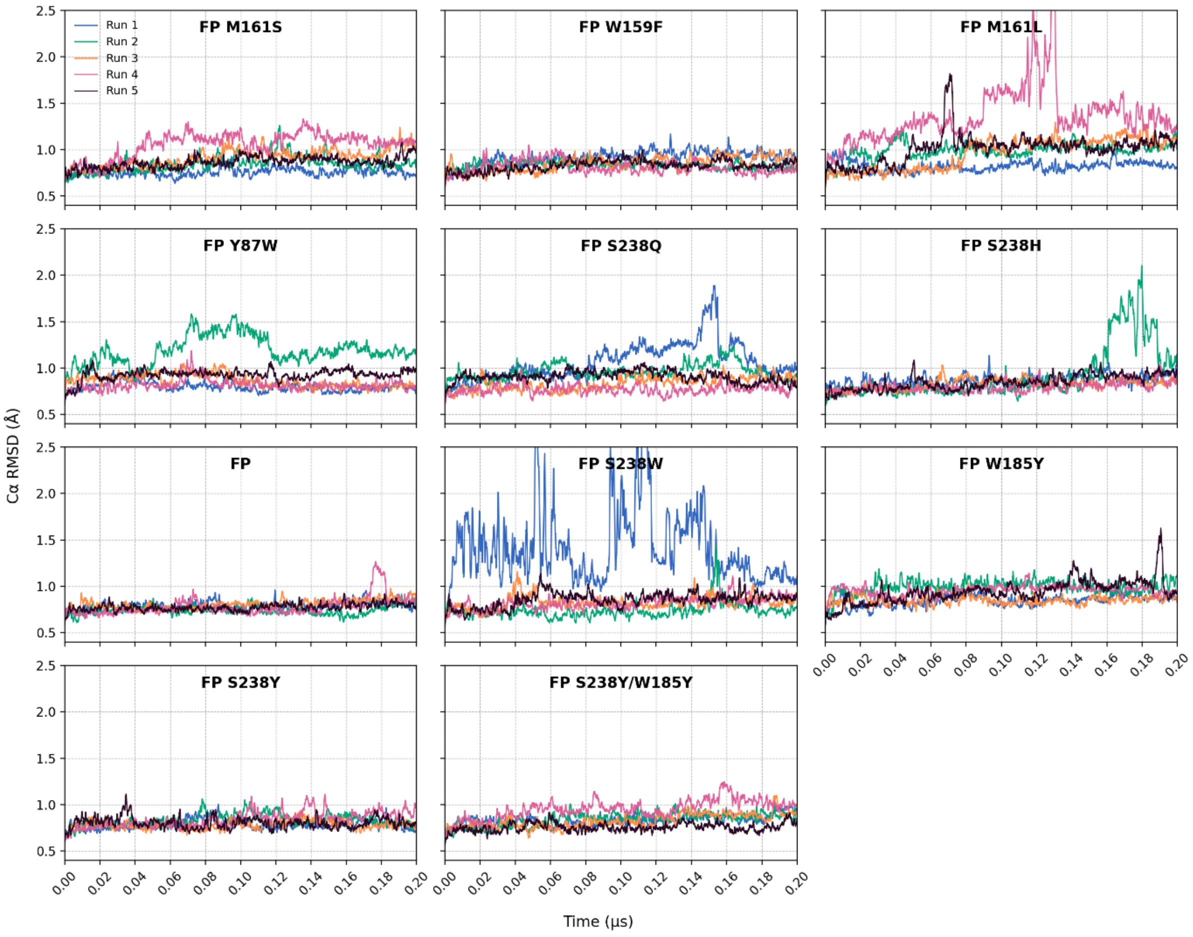

**Figure S8**. Root-mean-square deviation (RMSD) of Cα atoms for each mutant across five independent molecular dynamics (MD) simulations. Each coloured line represents one replicate run. Plots are ordered by in vitro fold change, from lowest to highest.

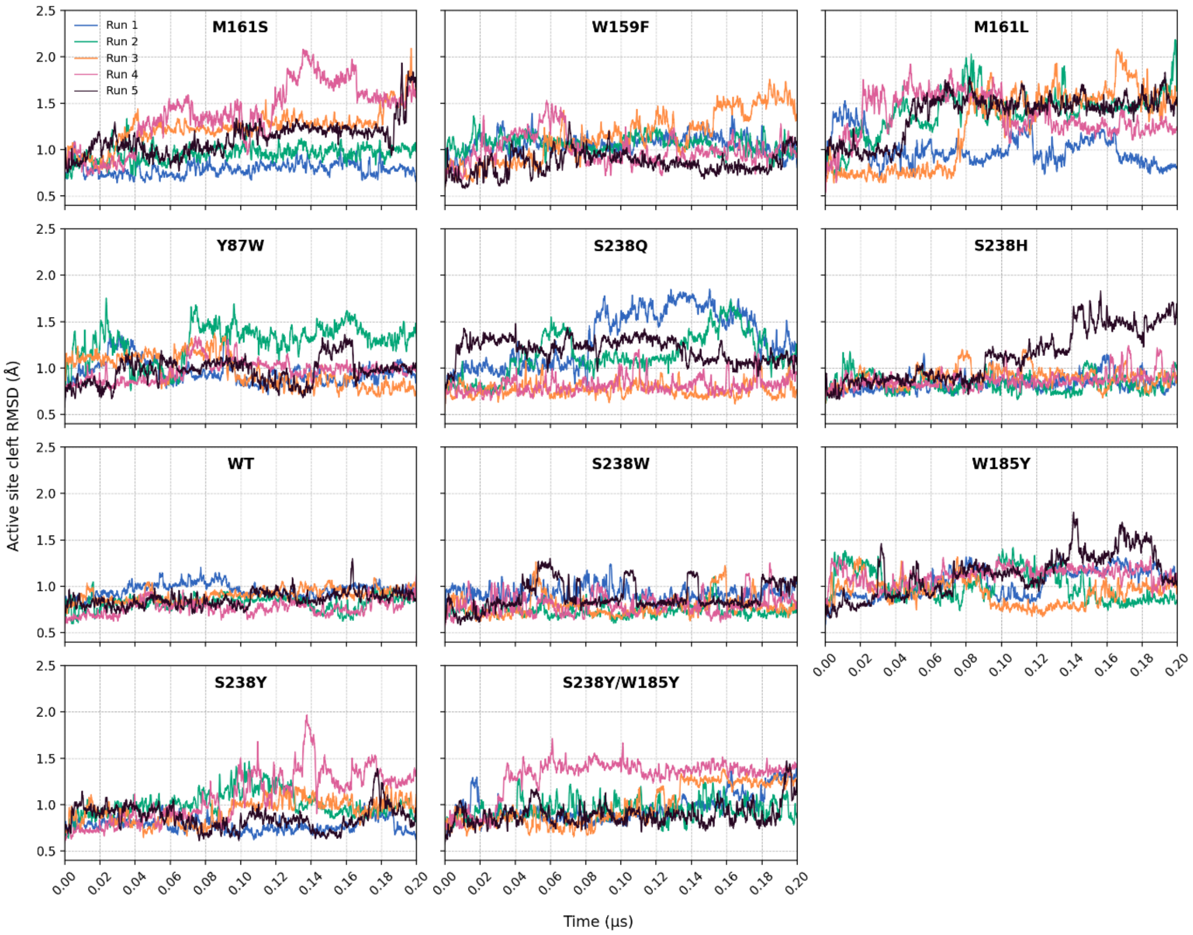

**Figure S9**. Root-mean-square deviation (RMSD) of the active site cleft residues for each mutant across five independent molecular dynamics (MD) simulations and fitted to the backbone of to the first frame of each trajectory. Each coloured line represents one replicate run. Plots are ordered by in vitro fold change, from lowest to highest.

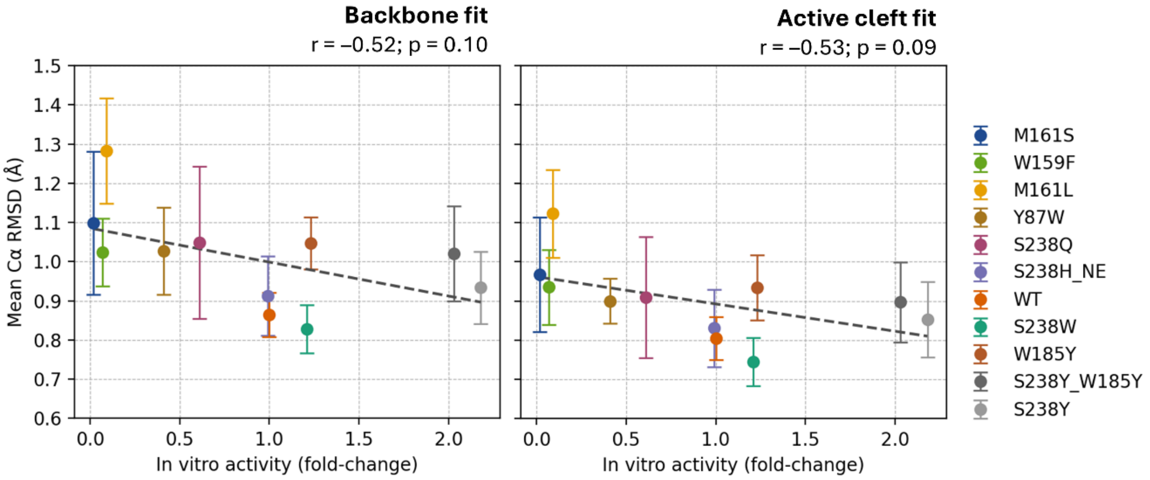

**Figure S10.** Relationship between in vitro activity and mean active site cleft Cα RMSD for each PETase mutant. Mean RMSD values were calculated from five replicate simulations after backbone fitting (left) or active-site cleft fitting (right), with error bars representing 95% confidence intervals (± 2σ) from 1000 bootstrap resampling iterations. Dashed lines indicate least-squares linear fits (Backbone fit: $y = -0.09x + 1.09$, $r = -0.52$, $p = 0.10$; Active cleft fit: $y = -0.07x + 0.96$, $r = -0.53$, $p = 0.09$). No statistically significant correlations were observed ($p > 0.05$).

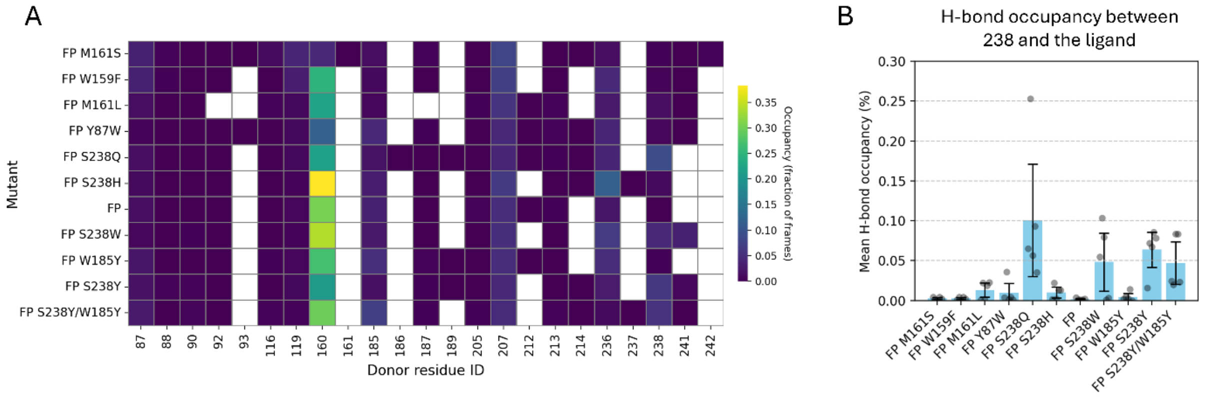

**Figure S11**. Analysis of hydrogen-bond formation between the ligand and the active site cleft residues occurring during the molecular dynamics (MD) simulations performed for the FP variants. (A) Heatmap of per-donor residue hydrogen bond occupancy between the ligand and the active site cleft residues in the FP mutants. Colours represent the fraction of MD simulation frames in which a hydrogen bond is observed for a given donor residue. White squares indicate no observed hydrogen bond in any frame. (B) Mean hydrogen bond occupancy per mutant for donor residue 238-ligand pair. Bars show the bootstrap mean ± 95 % confidence interval. Grey dots represent the values from five independent MD replicates used for the bootstrap calculation.

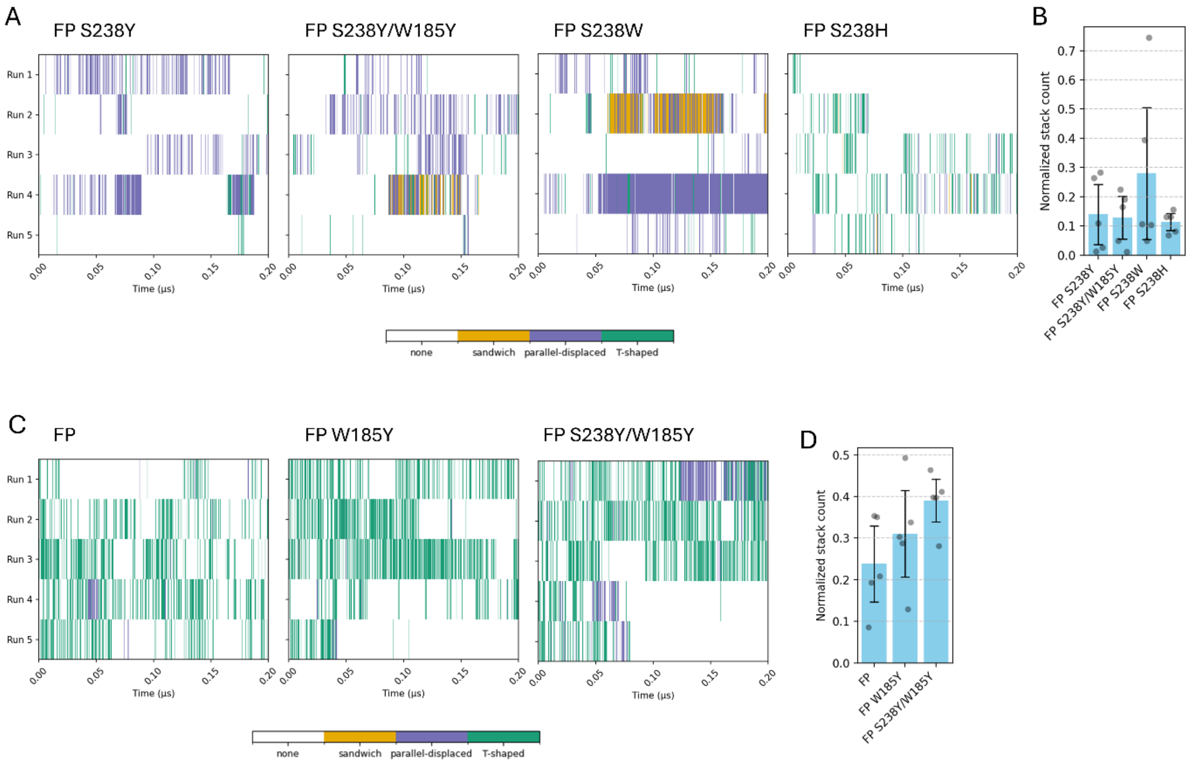

**Figure S12**. Analysis of relevant aromatic interactions occurring during the molecular dynamics (MD) simulations performed for the FP variants. **(A)** Time-resolved π-stacking interactions between residue 238 and the ligand for each MD replicate. Heatmaps show the assigned π-stacking type for each frame of each replicate simulation. **(B)** Mean normalized aromatic/stacking interactions (fraction of bound frames in which π-π stacking occurs divided by the total number of bound frames) between each residue in position 238 and the ligand. Bars show the bootstrap mean ± 95% confidence interval. Grey dots represent the values from five independent MD replicates used for the bootstrap calculation. **(C)** Time-resolved π-stacking interactions between residue 185 and the ligand for each MD replicate. Heatmaps show the assigned π-stacking type for each frame of each replicate simulation. **(D)** Mean normalized aromatic/stacking interactions (fraction of bound frames in which π-π stacking occurs divided by the total number of bound frames) between each residue in position 185 and the ligand. Bars show the bootstrap mean ± 95 % confidence interval. Grey dots represent the values from five independent MD replicates used for the bootstrap calculation.

### Supporting Tables

**Table S1**. List of prohibited amino acid substitutions for each of the 12 targeted amino acid residues at the active site. The amino acid residues were grouped according to their properties: positive residues (Arg, His, and Lys), negative residues (Asp and Glu), polar residues (Gln, Asn, Ser, Thr), non-polar residues (Ala, Val, Ile, Leu, Met, Phe, Tyr, and Trp), and special residues (Gly and Pro).

| **Residue position** | **Prohibited substitutions** |
|---|---|
| Y87 | Positive and special |
| T88 | Positive, special and Phe, Trp and Tyr |
| R90 | Positive, special and non-polar |
| S92 | Positive, special and non-polar except Ala |
| S93 | Positive, special and Met, Phe, Trp and Tyr |
| Q119 | Positive except Lys, Pro and non-polar |
| W159 | Positive except Lys, special and polar |
| M161 | Positive and special |
| W185 | Positive and special |
| I208 | Positive, special and Met, Phe, Trp and Tyr |
| S238 | Positive and special |
| N241 | Positive and special |

**Table S2.** List of PET hydrolase sequences retrieved from the PET-PAZy database and ordered according to the percentage identity, as calculated using the BlastP. Microbial host and enzyme annotations are presented as listed in the PET-PAZy database. GenBank or UniProt accession numbers are shown.

| **Microbial host, enzyme, gene** | **GenBank/ UniProt entry** | **Percentage Ident (%)** | **Length** |
|---|---|---|---|
| *Ideonella sakaiensis* 201-F61 , ISPETase, ISF6_4831 | A0A0K8P6T7 | 100 | 290 |
| *Acidovorax delafieldii*, PbsA/AdCut | Q8RR62 | 82.06 | 304 |
| *Rhizobacter gummiphilus* NS21, RgPETase/RgCut-I | A0A1W6L588 | 81.3 | 292 |
| *Caldimonas brevitalea*, PET12 (PbCut;SbCut), AAW51_2473 | A0A0G3BI90_9BURK | 66.79 | 298 |
| *Rhizobacter gummiphilus*, RgCut-II | WP_085749238.1 | 65.62 | 279 |
| *Burkholderia bacterium*, BurPL | A0A1F4JXW8 | 65.23 | 420 |
| *Oleispira antarctica* RB-8, PET5, LipA (=Oacut) | R4YKL9 | 54.83 | 310 |
| *T. cellulosilytica*, Enzyme 711 | WP_083947829.1 | 53.58 | 284 |
| Caldimonas taiwanensis+D57, CtPL (Enzyme 504) | WP_062195544.1 | 53.38 | 292 |
| *Marinobacter* sp., PLE629 | UUT36763.1 | 53.23 | 316 |
| Metagenome-derived, no obvious affiliation, PET2, lipIAF5-2 | C3RYL0 | 53.21 | 308 |
| *T. alba* DSM43185, Tha_Cut1, cut1 | E9LVH7 | 53.2 | 262 |
| *Ketobacter* sp., Enzyme 403 | RLU00646.1 | 53.08 | 312 |
| *Kibdelosporangium* aridum, KaPETase | A0A1W2FU08 | 52.42 | 286 |
| *Pseudomonas oleovorans/ pseudoalcaligenes* DSM 50188, PpCutA/PoCut | ADK73612.1 | 52.14 | 302 |
| *Deinococcus maricopensis* DSM 21211, PET1/DmPETase | E8U721 | 52.08 | 315 |
| *Halopseudomonas bauzanensis*, PbauzCut | A0A031MKR8 | 51.8 | 302 |
| *T. cellulosilytica* DSM44535, Thc_Cut1 | ADV92526.1 | 51.71 | 262 |
| *Pseudomonas paralcaligenes* MRCP1333, PpPETase | WP_220815207.1 | 51.57 | 298 |

| | | | |
|---|---|---|---|
| *Streptomyces calvus* DSM 41452, ScPETase | QDI72884.1 | 51.3 | 300 |
| *T. fusca* NRRL B-8184, Cut2 | AET05799.1 | 51.11 | 301 |
| *T. fusca* (strain YX), WSH03-11, Tfu_0883 | Q47RJ6 | 51.11 | 301 |
| *T. fusca,* TfCut_2 (Cut-2.kw3) | Q6A0I4 | 51.11 | 301 |
| *T. fusca,* TfCut_1 (Cut-1.kw3) (only active on 3PET; not on higher polymers!) | E5BBQ2 | 50.74 | 319 |
| *Marinobacter* sp., PLE628 | UUT36764.1 | 50.58 | 307 |
| *Actinobacteria bacterium* OV320, Enzyme 607 | WP_107095481.1 | 50.53 | 310 |
| *T. fusca,* Enzyme 701 | WP_104613137.1 | 50.37 | 301 |
| *Pseudomonas saudimassiliensis,* PsCut | A0A078MGG8 | 50.34 | 302 |
| *T. cellulosilytica* DSM44535, Thc_Cut2 | ADV92527.1 | 50.19 | 262 |
| *Amycolatopsis* bacterium, PET40 | WAU86704.1 | 50.19 | 263 |
| *T. alba (*AHK119*),* Est1 (Hydrolase 4); Enzyme 708 | BAI99230.2 | 50.19 | 296 |
| *Comamonadaceae bacterium*, Enzyme 406 | ODU60407.1 | 50 | 305 |
| *T. fusca* NRRL B-8184, Cut1 | AET05798.1 | 49.63 | 319 |
| *T. fusca* (strain YX), WSH03-11, Tfu_0882 | Q47RJ7 | 49.63 | 319 |
| *T. alba* AHK119, Est119, est2 | F7IX06 | 49.55 | 240 |
| *T. halotolerans,* Thh_Est | H6WX58 | 49.43 | 262 |
| Compost metagenome, PHL-5[2] | SAY37587.1 | 49.23 | 257 |
| *Pseudomonas alcaligenes*, PaCut | SUD16364.1 | 49.13 | 298 |
| *Pseudomonas pelagia* DSM 25163, PpelaLip | ANP21910.1 | 49.06 | 307 |
| *T. curvata* DSM43183, Tcur0390 | D1A2H1 | 49.06 | 292 |
| LCC, leaf compost metagenome, highly similar to HRB29 locus GBD22443 | G9BY57 | 48.89 | 293 |
| Compost metagenome, PHL-3[2] | SAY37583.1 | 48.86 | 259 |
| *Halopseudomonas formosensis, Hfor*_PE-H | WP_090538641.1 | 48.84 | 304 |
| *Microbispora sp.*, SIBER-1 | WOR09923.1 | 48.84 | 257 |
| *T. curvata* DSM43183, Tcur_1278 | D1A9G5 | 48.65 | 289 |
| *Vibrio gazogenes,* PET6, *BSQ33_03270* | A0A1Z2SIQ1 | 48.41 | 298 |
| *Thermoanaerobacter* sp. , PHL-7 (PES-H1; PES-H2) | SAY37592.1 | 48.28 | 259 |
| Compost metagenome, PHL-6[2] | SAY37589.1 | 48.08 | 257 |
| Compost metagenome, PHL-4[2] | SAY37584.1 | 48.08 | 257 |

| | | | |
|---|---|---|---|
| Compost metagenome, PHL-1[2] | SAY37579.1 | 47.89 | 260 |
| *Halopseudomonas aest*usnigri VGXO14, PE-H, *B7O88_11480* | 00089F5A9D | 47.87 | 304 |
| Compost metagenome, PHL-2[2] | SAY37582.1 | 47.71 | 260 |
| BhrPETase from HR29 bacterium, >96% identical to LCC | GBD22443.1 | 46.67 | 293 |
| *Saccharomonospora (Thermoactinomyces) viridis* AHK190, Cut190 | W0TJ64 | 45.49 | 304 |
| *Saccharopolyspora flava,* Enzyme 611 | WP_093412886.1 | 45.3 | 293 |
| *Moraxella* sp.TA144, lip1, Mors1 | P19833 | 45.24 | 319 |
| *Marinactinospora thermotolerans,* MtCut / Enzyme 606 | WP_078759821.1 | 44.48 | 311 |
| *Nocardioidaceae bacterium,* Enzyme 503 | EGD44994.1 | 43.53 | 294 |
| *Cryptosporangium aurantiacum*, *Ca*PETase | SHM40309.1 | 42.96 | 299 |
| *Actinobacteria bacterium* OK074, Enzyme 405 | WP_082414832.1 | 42.42 | 302 |
| Geothermal metagenome, Sis | XDS72785.1 | 41.87 | 282 |
| *Streptomyces* sp. SM14, SM14est | DAC80635.1 | 41.38 | 284 |
| *Allorhizocoloa rhizosphaerae*, Enzyme 407 | WP_117215036.1 | 40.7 | 434 |
| *Kaistella* (*Chryseobacterium*) *jeonii*, PET30 | WP_039353427.1 | 38.49 | 366 |
| *Aequorivita* sp. CIP111184, PET27 | WP_111881932.1 | 38.46 | 364 |
| *Ketobacter sp.*, Enzyme 409 | RLT92980.1 | 35.6 | 269 |
| *Caldibacillus thermoamylovorans*, Ces19_14 | WP_034767800.1 | 35.42 | 240 |
| *Ketobacter alkanivorans,* Enzyme 412 | WP_101893509.1 | 34.8 | 283 |
| *Pseudomonadota* bacterium, dsPETase06 | MEC8523093.1 | 31.28 | 292 |
| *Pseudomonas mendocina* ATCC 53552, PmC | N20M5AZM016 | 28.06 | 258 |
| Candidatus *Bathyarchaeota archaeon*, PET46 | RLI42440.1 | 28 | 262 |
| *Brucella*, PD3 | QPA27412.1 | 24.19 | 352 |

**Table S3.** List of primers used for cloning in this study.

| **Primer name** | **Sequence (5’->3’)** |
|---|---|
| pETDuet_HistagNCS_fwd | agccaggatccgaattcg |
| pETDuet_HistagNCS_rev | gtggtgatgatggtgatg |
| IsPETase_noSP_Histag_fwd | gccatcaccatcatcaccacCAAACTAACCCGTACGCAC |
| IsPETase_Histag_rev | ctcgaattcggatcctggctTCAAGAGCAATTCGCGGTAC |
| DuetUp1 | ggatctcgacgctctccct |
| DuetDown1 | gattatgcggccgtgtacaa |

**Table S4.** List of primers used for mutagenesis in this study. Base pairs that introduce the respective mutations are shown in red.

| **Primer name** | **Sequence (5'->3')** |
|---|---|
| W159F_fwd | gtatgggggtgatgggctt**tc**tcgatgggtgg |
| W159F_rev | ccacccatcga**ga**agcccatcacccccatac |
| S238Y_fwd | cttgagattaagggaggatcccatt**at**tgtgccaattcgg |
| S238Y_rev | ccgaattggcaca**at**aatgggatcctcccttaatctcaag |
| S238Q_fwd | agattaagggaggatcccat**ca**gtgtgccaattcgggaaac |
| S238Q_rev | gtttcccgaattggcacac**tg**atgggatcctcccttaatct |
| S238W_fwd | agggaggatcccatt**g**gtgtgccaattcgg |
| S238W_rev | ccgaattggcacac**c**aatgggatcctccct |
| S238H_fwd | ttccttgagattaagggaggatcccat**cat**tgtgccaattcgggaaa |
| S238H_rev | tttcccgaattggcaca**atg**atgggatcctcccttaatctcaaggaa |
| W185Y_fwd | ctgcaccccaggcgcctt**at**cattcatcgaccaatttt |
| W185Y_rev | aaaattggtcgatgaatg**at**aaggcgcctggggtgcag |
| Y87W_fwd | ccattgcgatcgtaccaggat**gg**accgcccgcca |
| Y87W_rev | tggcgggcggt**cc**atcctggtacgatcgcaatgg |
| M161L_fwd | atgggctggtcg**t**tgggtgggggag |
| M161L_rev | ctcccccaccca**a**cgaccagcccat |
| M161S_fwd | gatgggctggtcga**gc**ggtgggggaggc |
| M161S_rev | gcctcccccacc**gc**tcgaccagcccatc |

**Table S5.** List of production strains used in this study.

| **Strain** | **Host** | **Relevant genotype** | **Source** |
|---|---|---|---|
| *E. coli* BL21 (DE3) | - | *F_ompT gal dcm lon hsdSB(rB- mB-) λ(DE3 *lacI lacUV5-T7 gene 1 ind1 sam7 nin5])* | NZYtech |
| PET1 | *E. coli* BL21 (DE3) | PETDuet_6xHistag_IsPETase_noSP | This study |
| PET3 | *E. coli* BL21 (DE3) | pETDuet_6xHisTag_IsPETase_S121E_D1866H_R280A_noSP | This study |
| PET4 | *E. coli* BL21 (DE3) | pETDuet_6xHisTag_IsPETase_S121E_D1866H_R280A_R224Q_N233K_noSP | This study |
| PET5 | *E. coli* BL21 (DE3) | pETDuet_6xHisTag_IsPETase_S121E_D1866H_R280A_R233C_S282C_noSP | This study |
| PET6 | *E. coli* BL21 (DE3) | pETDuet_6xHisTag_IsPETase_S121E_D1866H_R280A_R224Q_N233K_Y87W_noSP | This study |
| PET7 | *E. coli* BL21 (DE3) | pETDuet_6xHisTag_IsPETase_S121E_D1866H_R280A_R224Q_N233K_W159F_noSP | This study |
| PET8 | *E. coli* BL21 (DE3) | pETDuet_6xHisTag_IsPETase_S121E_D1866H_R280A_R224Q_N233K_M161L_noSP | This study |
| PET9 | *E. coli* BL21 (DE3) | pETDuet_6xHisTag_IsPETase_S121E_D1866H_R280A_R224Q_N233K_M161S_noSP | This study |
| PET10 | *E. coli* BL21 (DE3) | pETDuet_6xHisTag_IsPETase_S121E_D1866H_R280A_R224Q_N233K_S238Y_noSP | This study |
| PET11 | *E. coli* BL21 (DE3) | pETDuet_6xHisTag_IsPETase_S121E_D1866H_R280A_R224Q_N233K_S238W_noSP | This study |
| PET12 | *E. coli* BL21 (DE3) | pETDuet_6xHisTag_IsPETase_S121E_D1866H_R280A_R224Q_N233K_S238Q_noSP | This study |
| PET13 | *E. coli* BL21 (DE3) | pETDuet_6xHisTag_IsPETase_S121E_D1866H_R280A_R224Q_N233K_W185Y_noSP | This study |
| PET14* | *E. coli* BL21 (DE3) | pETDuet_6xHisTag_IsPETase_S121E_D1866H_R280A_R224Q_N233K_Y87W_M161S_noSP | This study |
| PET15* | *E. coli* BL21 (DE3) | pETDuet_6xHisTag_IsPETase_S121E_D1866H_R280A_R224Q_N233K_S238Y_M161S_noSP | This study |
| PET16 | *E. coli* BL21 (DE3) | pETDuet_6xHisTag_IsPETase_S121E_D1866H_R280A_R224Q_N233K_S238H_noSP | This study |
| PET17 | *E. coli* BL21 (DE3) | pETDuet_6xHisTag_IsPETase_S121E_D1866H_R280A_R224Q_N233K_S238Y_W185Y_noSP | This study |

| | | | |
|---|---|---|---|
| PET18 | *E. coli* BL21 (DE3) | pETDuet_6xHisTag_IsPETase_S121E_D1866H_R280A_R233C_S282C_S238Y_noSP | This study |
| PET19 | *E. coli* BL21 (DE3) | pETDuet_6xHisTag_IsPETase_S121E_D1866H_R280A_R233C_S282C_W185Y_noSP | This study |
| PET20 | *E. coli* BL21 (DE3) | pETDuet_6xHisTag_IsPETase_S121E_D1866H_R280A_R233C_S282C_S238Y_W185Y_noSP | This study |

* Two double mutants initially tested in vitro (FP Y87W/M161S and FP S238Y/M161S; strains PET14 and PET15, respectively) were selected from a preliminary PETase WT-based variant listing. Subsequent updates to the screening pipeline rendered theses designs less relevant and less catalytically promising, as they ranked much lower in the FP round. In addition, both exhibited negligible hydrolytic activity toward PET disks in vitro (< 0.1-fold change compared to FP). For these reasons, they were not included in the main analysis.

**Table S6.** Complete library of filtered and ranked single mutants from the GDEE platform. Mutations selected for in vitro validation are highlighted in red. FastPETase is highlighted in blue.

| | **Filtering by poses A and B** | | **Filtering by poses C and D** | |
|---|---|---|---|---|
| **Rank** | **Mutant** | **Energy (kcal mol$^{-1}$)** | **Mutant** | **Energy (kcal mol$^{-1}$)** |
| 1 | S92Q | −7.1 | **S238Y** | −7.3 |
| 2 | S92N | −7.1 | S238F | −7.2 |
| 3 | **S238Y** | −7.1 | **W159F** | −7 |
| 4 | **S238Q** | −7.1 | **S238W** | −7 |
| 5 | N241Q | −7.1 | S92Q | −6.9 |
| 6 | S238R | −7.1 | S92N | −6.9 |
| 7 | R90T | −7 | S92A | −6.9 |
| 8 | T88Q | −7 | S92C | −6.9 |
| 9 | S92T | −7 | S93N | −6.9 |
| 10 | S92A | −7 | S238L | −6.9 |
| 11 | S92C | −7 | **S238Q** | −6.9 |
| 12 | S93V | −7 | S238N | −6.9 |
| 13 | S93N | −7 | **Y87W** | −6.9 |
| 14 | **W159F** | −7 | N241Q | −6.9 |
| 15 | N241L | −7 | N241T | −6.9 |
| 16 | N241A | −7 | N241V | −6.9 |
| 17 | N241M | −7 | N241M | −6.9 |
| 18 | **S238H** | −7 | S238K | −6.9 |
| 19 | Q119G | −6.9 | S238R | −6.9 |
| 20 | S93A | −6.9 | **S238H** | −6.9 |
| 21 | R90C | −6.9 | Q119G | −6.8 |
| 22 | R90Q | −6.9 | S93A | −6.8 |
| 23 | R90S | −6.9 | R90C | −6.8 |
| 24 | Q119N | −6.9 | S92T | −6.8 |
| 25 | Q119S | −6.9 | Q119N | −6.8 |
| 26 | R90N | −6.9 | Q119S | −6.8 |
| 27 | S93C | −6.9 | S93T | −6.8 |
| 28 | S93I | −6.9 | S238T | −6.8 |

| | | | | |
|---|---|---|---|---|
| 29 | T88C | −6.9 | S238V | −6.8 |
| 30 | S93L | −6.9 | S238I | −6.8 |
| 31 | **M161S** | −6.9 | N241L | −6.8 |
| 32 | S238A | −6.9 | N241I | −6.8 |
| 33 | S238L | −6.9 | N241S | −6.8 |
| 34 | T88M | −6.9 | N241A | −6.8 |
| 35 | N241I | −6.9 | N241W | −6.8 |
| 36 | N241S | −6.9 | R90T | −6.7 |
| 37 | N241C | −6.9 | R90Q | −6.7 |
| 38 | N241V | −6.9 | T88N | −6.7 |
| 39 | N241W | −6.9 | R90S | −6.7 |
| 40 | S238K | −6.9 | S93V | −6.7 |
| 41 | T88I | −6.8 | R90N | −6.7 |
| 42 | T88A | −6.8 | Q119K | −6.7 |
| 43 | Q119K | −6.8 | S93C | −6.7 |
| 44 | Q119T | −6.8 | T88C | −6.7 |
| 45 | S238V | −6.8 | S93L | −6.7 |
| 46 | M161V | −6.8 | S93Q | −6.7 |
| 47 | **W185Y** | −6.8 | M161Q | −6.7 |
| 48 | S238F | −6.8 | S238A | −6.7 |
| 49 | S238I | −6.8 | W159H | −6.7 |
| 50 | S238N | −6.8 | FastPETase | −6.7 |
| 51 | T88V | −6.8 | I208L | −6.7 |
| 52 | N241T | −6.8 | S238C | −6.7 |
| 53 | N241Y | −6.8 | S238M | −6.7 |
| 54 | T88N | −6.7 | T88V | −6.7 |
| 55 | Q119C | −6.7 | T88L | −6.7 |
| 56 | S93T | −6.7 | T88M | −6.7 |
| 57 | S93Q | −6.7 | N241C | −6.7 |
| 58 | S238T | −6.7 | N241F | −6.7 |
| 59 | M161Q | −6.7 | T88Q | −6.6 |
| 60 | W159H | −6.7 | T88I | −6.6 |
| 61 | **M161L** | −6.7 | Q119C | −6.6 |
| 62 | W159Y | −6.7 | T88A | −6.6 |

| | | | | |
|---|---|---|---|---|
| 63 | FastPETase | −6.7 | Q119T | −6.6 |
| 64 | S238C | −6.7 | S93I | −6.6 |
| 65 | **S238W** | −6.7 | **M161S** | −6.6 |
| 66 | S238M | −6.7 | W159I | −6.6 |
| 67 | T88L | −6.7 | T88S | −6.6 |
| 68 | T88S | −6.7 | Y87N | −6.6 |
| 69 | **Y87W** | −6.7 | N241Y | −6.6 |
| 70 | Y87T | −6.7 | **W185Y** | −6.5 |
| 71 | Y87Q | −6.7 | I208S | −6.5 |
| 72 | W159L | −6.6 | I208N | −6.5 |
| 73 | W159M | −6.6 | I208V | −6.5 |
| 74 | I208T | −6.6 | Y87F | −6.5 |
| 75 | N241F | −6.6 | Y87T | −6.5 |
| 76 | M161C | −6.5 | Y87Q | −6.5 |
| 77 | Y87A | −6.5 | Y87L | −6.5 |
| 78 | Y87F | −6.5 | Y87M | −6.5 |
| 79 | Y87S | −6.5 | Y87A | −6.4 |
| 80 | Y87N | −6.5 | W185V | −6.4 |
| 81 | W159I | −6.4 | W159L | −6.4 |
| 82 | I208V | −6.4 | W159Y | −6.4 |
| 83 | Y87C | −6.4 | I208T | −6.4 |
| 84 | W159C | −6.3 | W185T | −6.3 |
| 85 | W159V | −6.3 | M161A | −6.3 |
| 86 | M161I | −6.3 | M161I | −6.3 |
| 87 | I208N | −6.3 | I208C | −6.3 |
| 88 | I208Q | −6.3 | W159M | −6.3 |
| 89 | Y87L | −6.3 | Y87C | −6.3 |
| 90 | Y87M | −6.3 | W159C | −6.2 |
| 91 | I208C | −6.2 | W185N | −6.2 |
| 92 | I208L | −6.2 | I208Q | −6.2 |
| 93 | I208S | −6.1 | Y87S | −6.2 |
| 94 | M161A | −6 | W159V | −6.1 |
| 95 | W185F | −5.9 | M161C | −6 |
| 96 | W185T | −5.8 | M161V | −6 |
| 97 | W185V | −5.8 | **M161L** | −6 |

| | | | | |
|---|---|---|---|---|
| 98 | W185A | −5.5 | W185F | −5.8 |
| 99 | W185N | −5.5 | W185S | −5.7 |
| 100 | M161N | −5.3 | W185A | −5.7 |
| 101 | M161T | −5.3 | W185I | −5.6 |
| 102 | W159A | −5.2 | W185C | −5.6 |
| 103 | I208A | −5.2 | W185M | −5.5 |
| 104 | W185S | −5 | I208A | −5.4 |
| 105 | W185I | −5 | M161N | −5.2 |
| 106 | W185C | −5 | M161T | −5.2 |
| 107 | | | W159A | −5 |
| 108 | | | W185L | −4.4 |

**Table S7**. List of mutations and their respective binding free energies, selected from the filtered and ranked poses from the GDEE library, along with FastPETase variant shown as reference.

| **Mutation** | **Binding free energy (kcal mol$^{-1}$)** | |
|---|---|---|
| | **Filtering by poses' I and II rules** | **Filtering by poses' III and IV rules** |
| S238Y | −7.1 | −7.3 |
| S238H | −7.1 | −6.9 |
| S238Q | −7.1 | −6.9 |
| W159F | −7.0 | −7.0 |
| M161S | −6.9 | −6.6 |
| FastPETase | −6.9 | −6.7 |
| W185Y | −6.8 | −6.5 |
| S238W | −6.7 | −7.0 |
| Y87W | −6.7 | −6.0 |
| M161L | −6.7 | −6.9 |
